# A critical role for the ER Membrane Complex (EMC) in lipid droplet homeostasis

**DOI:** 10.1101/2025.03.10.642408

**Authors:** Érika de Carvalho, Abdulbasit Amin, Richard Little, Keily Fonseca Silva, Catarina Gaspar, Marina Badenes, Emma Burbridge, Tiago Paixão, Maria João Amorim, Pedro Faísca, Roman Fischer, Iolanda Vendrell, Darragh P. O’Brien, Kvido Strisovsky, John C. Christianson, Pedro M. Domingos, Colin Adrain

## Abstract

The ER Membrane complex (EMC) facilitates insertion of multiple classes of integral membrane protein into the lipid bilayer of the endoplasmic reticulum (ER). Recent years have seen significant progress in understanding the structural aspects of EMC function, but the client protein/s enforcing its evolutionary conservation throughout eukaryotic kingdoms remain to be determined. Given reported role/s for the EMC in lipid homeostasis, we deleted the essential EMC3 subunit in the liver and adipose tissues of mice and examined the impact of EMC loss in a range of metabolic models. Intriguingly, absence of EMC3 in either the liver or adipose tissues results in defective storage of triglycerides and perturbations in lipid droplet (LD) homeostasis. EMC-deficient mouse adipose tissues were defective in their ability to mediate fat storage in response to a high fat diet and were unable to support non-shivering thermogenesis. Proteomic analyses of EMC3-deficient tissues found reduced levels of FITM2, a key regulator of LD biogenesis and ER homeostasis, identifying it as a novel EMC client. Strikingly, deletion of the EMC3 homolog *dPob* in the *Drosophila* fat body, which shares overlapping functions with mammalian liver/adipose tissues, revealed similar defects in LD homeostasis. Together, our results indicate that the EMC plays a key evolutionarily conserved role in biogenesis of machinery maintaining triglyceride storage in metazoans.

## Introduction

Mammalian genomes encode approximately 5000 integral membrane proteins that fulfil fundamental roles in biology, including signalling, lipid biogenesis/trafficking and the transport of ions and nutrients. Most integral membrane proteins are inserted into the lipid bilayer at the endoplasmic reticulum (ER), via their transmembrane domains (TMDs), alpha helical segments enriched in amino acids with hydrophobic side chains^1^.

Integral membrane proteins are inserted into the ER lipid bilayer by molecular assemblages called “insertases”^1^. During this process, the insertion machinery also interprets molecular cues that ensure the correct orientation of the protein’s N- and C-termini relative to the cytoplasm^1^. This process is more complex for polytopic membrane proteins, since multiple TMDs must be handled, in addition to loops of varying size between TMDs^1^.

The mechanisms of transmembrane protein insertion into the ER membrane have been studied extensively and it is now apparent that multiple insertion pathways are available. Phylogenetic, biochemical and structural studies indicate an evolutionarily conserved family of “insertase” molecules of the Oxa1 family^2,3^. This includes Sec61 (the first ER “insertase” to be identified); the GET complex insertase Get1, TMCO1, and EMC3, the insertase component of a multi-subunit assembly called the ER membrane complex (EMC)^3–5^.

Although Sec61 is long-established as the principal insertase, recent evidence has raised questions about its importance in determining polytopic membrane protein topogenesis. Interestingly, the EMC has emerged as an important point of triage for coordinated insertion of polytopic TMD containing proteins^6–8^. For instance, EMC enforces the correct topogenesis of the first TMD of certain G protein coupled receptors (GPCRs) that contain a signal anchor sequence^6^, while the remaining TMDs are inserted in an EMC-independent manner. In a similar vein, EMC inserts the “terminal” TMD of polytopic proteins that culminate in short C-terminal luminal tails^8^ where proximity between the TMD and C-terminus necessitates post-translational insertion^8^. In addition, EMC acts as the insertase for Tail-Anchored (TA) proteins, single-pass transmembrane proteins with a very short C-terminal tail immediately following their TMD, with moderate to low TMD hydrophobicity, that cannot be handled by the GET/TRC complex^9–11^, and some type III proteins^12^.

While much is known about the mechanistic roles of EMC, animal studies are required to identify the relevant endogenous EMC client proteins and to determine whether the EMC-dependency of a given client protein passes a threshold that renders the EMC essential for a given biological process *in vivo*. This has been hampered by the lethality associated with the global deletion of core EMC subunits that lead to the depletion of the entire EMC^10^,but are now emerging from tissue-specific KO studies. Studies in *Drosophila*^13–15^ and mice^15–17^ indicate that the EMC plays a key role in retinal development and homeostasis. In *Drosophila*, rhodopsin-1 biogenesis indirectly requires EMC, since the direct client of EMC is the TA protein Xport-A^11^, a chaperone that mediates rhodopsin-1 biogenesis by stabilizing its first five transmembrane domains ^18^.

Mouse KO studies have revealed an important role for the EMC in lung development, in the processing and routing of surfactant and for biogenesis of ABCA3, a key regulator of phospholipid transport into lamellar bodies, a secretory organelle where lung surfactants are produced^19^. Deletion of EMC3 in intestinal epithelial cells results in defects in intestinal barrier formation and is required for the stem cell niche function of paneth cells^20^. In contrast, ablation in mice of the non-core subunit EMC10 is viable, but associated with sperm dysfunction, mild behavioural defects, and delayed angiogenesis^21–23^. The EMC also plays critical roles in viral infection and is required for the biogenesis of key *Flavivirus* proteins^24,25^.

Multiple lines of evidence also implicate the EMC in the maintenance of lipid homeostasis. As noted above, the EMC is implicated in phospholipid transport in the lung^19^. In earlier studies in *Saccharomyces cerevisiae,* EMC was found to mediate phospholipid transfer between the ER and mitochondria^26^ while it is required for the biosynthesis of phosphatidylcholine (PC) or phosphatidylethanolamine (PE) in *Trypanosoma brucei*^27^. Studies using mammalian cells have uncovered a role for the EMC in sterol biogenesis (via the insertion of squalene synthase (SQS, also called FDFT1)) and esterification (via the biogenesis of enzymes sterol-O-acyltransferase 1 (SOAT1)^10^. Another EMC client protein, TMEM97 ^10,28^, negatively regulates the abundance of the intracellular cholesterol trafficking protein NPC1 (Niemann-Pick disease, type C1) and consequently, transfer of cholesterol to the ER^29^.

Given the strong link between EMC-mediated insertion and lipid homeostasis, we sought to explore this role in greater detail by knocking out EMC3 in murine liver and adipose tissues–key organs for lipid homeostasis^30,31^. Here we report an important evolutionarily conserved role for the EMC in lipid droplet homeostasis in metazoans and identify FITM2, a known mediator of LD biogenesis/homeostasis as a novel EMC-dependent client protein *in vivo*.

## Results

### Liver-specific ablation of *Emc3* results in hepatocyte death, compensatory proliferation and transient liver dysfunction

To ascertain the impact of EMC-dependent insertion on metabolic regulation and in particular, lipid homeostasis, we crossed mice harbouring a floxed allele of the EMC insertase subunit *Emc3* with animals expressing Albumin-cre, to ablate expression of *Emc3*, an essential EMC subunit, in hepatocytes. As expected, EMC3 was specifically ablated in the liver, but not in adipose tissue depots (**Fig.1a,b; Fig S1a**) from control floxed *Emc3^fl/fl^*(here and subsequently denoted as “WT”) versus mice with liver-specific deletion of *Emc3* (Emc ^Alb-cre Δ/Δ^, hence denoted “LKO”). As has been reported in other models ^10^, EMC3 depletion triggered loss of the other key EMC subunits (e.g. EMC5 and EMC6), precluding EMC complex assembly that leads to a loss of its functionality (**Fig. S1b,c**). Notably, EMC3-LKOs were born at the expected mendelian ratios (**Table 1**) and appeared ostensibly normal in terms of body size and gross appearance of the liver (**Fig.1c)**. These animals also appeared normal across several parameters including body weight, liver weight, and food intake (**Fig.1d-f**).

**Table 1.** Mendelian Ratios resulting from intercrosses between *Alb*^cre^ *Emc3*^△/△^ males or *Adipoq*^cre^ *Emc3*^△/△^ males and *Emc ^floxed^* females.

| Genotype | Number of animals (%) |  |
| --- | --- | --- |
|  | Males | Females |
| *Emc3<sup>lox/lox</sup>* (WT) | 73 (24.3%) | 64 (21.3%) |
| *Alb<sup>cre</sup> Emc3$\Delta/\Delta$* (KO) | 77 (25.7%) | 86 (28.7%) |
|  | Males | Females |
| *Emc3<sup>lox/lox</sup>* (WT) | 76 (25.3%) | 78 (26%) |
| *Adipoq<sup>cre</sup> Emc3$\Delta/\Delta$* (KO) | 74 (24.7%) | 72 (24%) |

\* A total of 300 animals were evaluated

Upon closer histopathological examination of LKO mouse livers at 8 weeks of age however, H&E staining revealed evidence of cell death and accompanying compensatory proliferation (**Fig.S1d**). This was confirmed by IHC staining for caspase-3 and Ki67 respectively (**Fig.1g-h**). Consistent with these histological observations, EMC3 LKO animals exhibited liver damage and hepatocyte death, reflected by significantly elevated levels of Alanine Aminotransferase (ALT) and Aspartate Aminotransferase (AST) in serum (**Fig.1i,j**). LKOs also showed a significant reduction in hepatic clearance of indocyanine green - a fluorescent dye which is normally rapidly cleared from circulation via the hepatic-biliary system following intravenous injection^32^ (**Fig. 1k**). This indicates that LKO animals also present with liver dysfunction. Notably, the EMC3-associated liver damage phenotype was most pronounced at 8 weeks of age, but receded with time, losing significance by the time animals reached 20 weeks of age (**Fig. 1l**). Importantly, we observed that EMC3 expression in LKO tissues had recovered to levels equivalent to that of WT by the time the animals had reached 33 weeks of age (**Fig. S1e**). Taking this into account, all subsequent experiments on liver-specific *Emc3* mutants were carried out in animals at 8-12 weeks of age. Overall, we interpret the transient liver phenotype to mean that the absence of EMC functionality causes hepatic dysfunction and hepatocyte death. The transient nature of phenotype could be attributable to mosaicism, which allows for compensatory proliferation of WT cells that have not undergone albumin-cre-mediated recombination, which has been observed in other liver-specific KO models ^33,34^ and other cre lines^35^.

**Figure 1.**
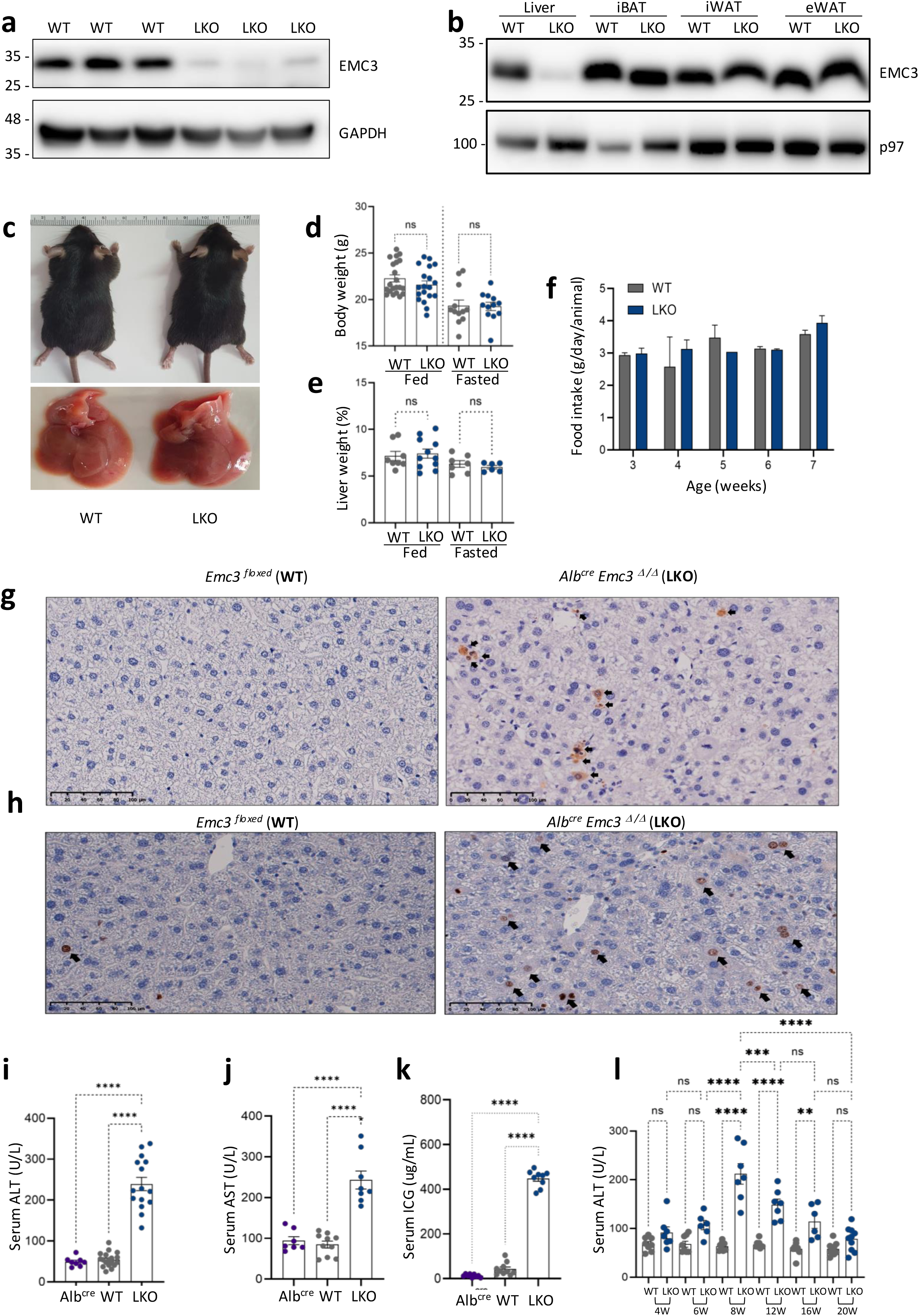
EMC3 KO liver-specific deletion (LKO) is viable but exhibit signs of liver damage and liver disfunction. **(a)** Absence of EMC3 protein in the liver of LKO animals was confirmed by western blotting with the indicated antibodies. A GAPDH probing is used as a loading control (**b)** Western blot panel corroborating the liver-specific deletion of EMC3 when compared against adipose tissue samples from the same animals. A p97 immunoblot is used as a loading control. **(c)** Compared to WT animals, LKO animals do not differ in their body appearance nor in their liver morphology. No significant difference is observed between the groups in their **(d)** body weight or **(e)** liver weight under *ad libitum* feeding (“fed”) or overnight fasting (“fasted”). **(f)** No difference is observed in food intake. **(g-h)** LKO animals exhibit a higher number of active caspase-3-positive cells compared to the WT controls. Images were obtained at 20X magnification. LKO animals present high serum levels of **(i)** alanine aminotransferase (ALT) and **(j)** aspartate aminotransferase (AST). **(k)** LKO animals present a high concentration of Indocyanine green (ICG) in the serum samples. Statistical analysis: One-way ANOVA – Tukey’s multiple comparison test. **(l)** LKO animals exhibit a decrease in serum ALT levels over time, indicating the recovery from liver damage. Data are from, at least, two distinct biological replicate experiments each containing N=3 mice per genotype. Each data point represents an individual animal; the same animals were compared at different time points throughout the experiment. Statistical analyses: one-way ANOVA – Tukey’s multiple comparison test. The error bars represent the standard error of the mean (SEM).ns, non-significant (p>0.05); **p< 0.01; ***p<0.001; ****p <0.0001.

Given the prior implication of the EMC in lipid biogenesis and trafficking in cellular studies^9,10,26,36^ and the liver’s crucial roles in lipid homeostasis, we next determined how EMC absence impacted various metabolic parameters. Consistent with previous findings from cells^9,10^, hepatic ablation of EMC3 substantially reduced levels of the EMC client squalene synthase (SQS) (**Fig.2a**). SQS mediates the first committed step in *de novo* sterol biogenesis, catalysing the conversion of the isoprenoid precursor farnesyl-pyrophosphate into squalene, which is then further modified in subsequent steps to produce cholesterol. Notably, hepatic deletion of *Emc3* did not significantly impact on serum (**Fig.2b**) or hepatic (**Fig.2c**) cholesterol levels. One possibility is that SQS depletion in our EMC3-deficient model (which as noted above may be prone to mosaicism and counter-selection for WT hepatocytes), does not reach the threshold where it impinges on sterol biogenesis. Consistent with this, SQS is not a rate-limiting enzyme in cholesterol biogenesis, since *Sqs* heterozygous mice exhibit normal hepatic cholesterol synthesis and normal plasma cholesterol levels^37^.

### Hepatic deletion of EMC3 causes defects in triglyceride homeostasis and LD defects

We next examined the impact of EMC3 ablation on triglyceride homeostasis, under standard chow feeding versus prolonged (12h) fasting conditions. Prolonged fasting drives a variety of metabolic adaptations including hepatic lipid droplet accumulation in response to elevated lipolysis in adipose tissues^38–40^. While the fed and fasted serum triglyceride (**Fig2d**) and fasted serum free glycerol (**Fig.2e**) were insensitive to hepatic *Emc3* deletion, the fasting-induced accumulation of triglycerides was significantly blunted in the KO animals compared to controls (**Fig.2f**). The observation that fasting hepatic levels but not serum levels are affected by ablation of EMC3 is consistent with the fact that the liver only stores a minor proportion (some 5%) of total body triglycerides^30^. As lipid droplets (LD) are the major cellular store of triglycerides, we used electron microscopy to inspect hepatocyte LDs following cardiac puncture, perfusion and fixation of livers from fasted mice. Intriguingly, the LDs of KO hepatocytes were much less prominent and less numerous (**Fig. 2g,h**) than control animals, suggesting a defect in LD accumulation or turnover in EMC3-deficient hepatocytes.

**Figure 2.**
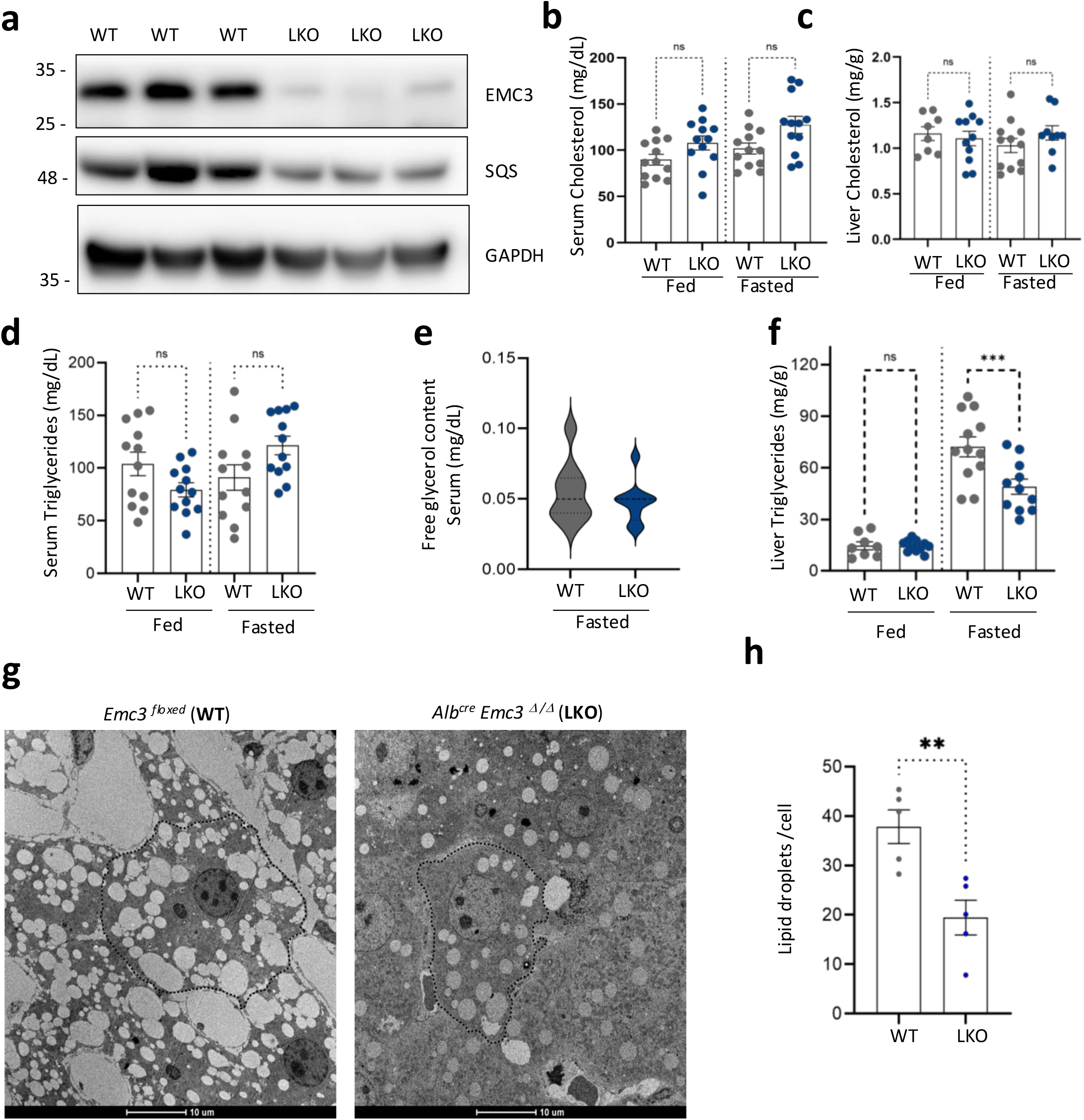
Hepatic deletion of EMC3 affects triglyceride and lipid droplet homeostasis. **(a)** Immunoblots indicating reduced expression of SQS in the liver of LKO animals compared to the WT controls. LKO animals show no significant difference in serum **(b)** and **(c)** hepatic cholesterol levels, independent of feeding conditions (fed or fasted). **(d,e)** No statistical difference in triglyceride or free glycerol levels was observed in serum samples. **(f)** LKO animals exhibit reduced levels of triglycerides in the liver after fasting. Statistical analysis: Ordinary One-way ANOVA – Šidák’s multiple comparison test. **(g)** Electron microscopy images demonstrating substantially reduced lipid droplet particles in the liver of LKO animals compared to WT controls, after fasting. **(h)** The mean value of number of lipid droplets per hepatocyte was enumerated. Each data point represents liver images sampled from an individual animal (5 per group). Statistical analysis: Shapiro-Wilk normal distribution, Outliers ROUT, and Unpaired Student t-test. Data are from three independent experiments (N=3 per genotype). Each data point represents an individual animal. The error bars represent the standard error of the mean (SEM). ns, non-significant (p>0.05); *p<0.05; **p <0.01 and ***p<0.001.

**Figure 3.**
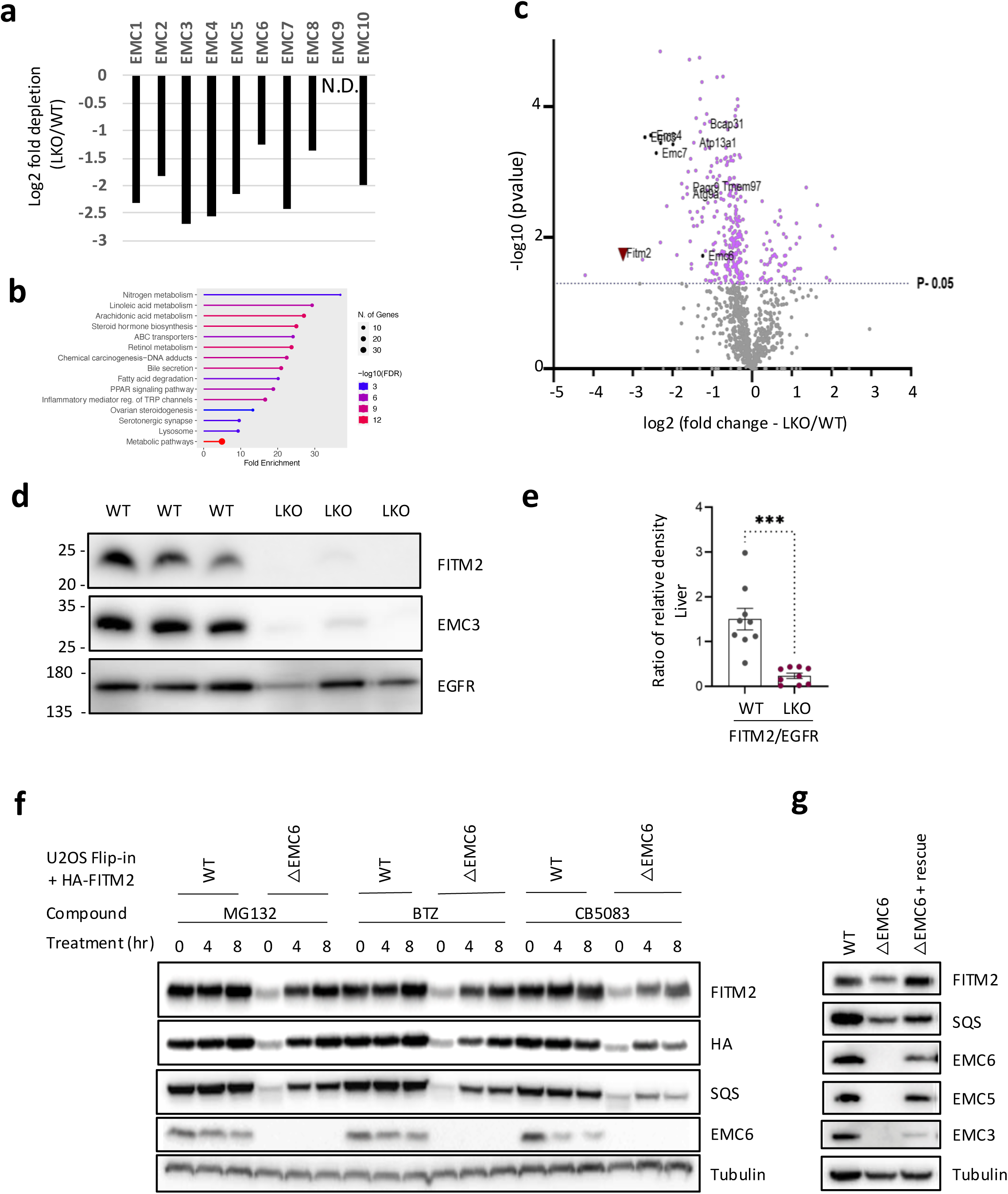
EMC3 absence results in reduced FITM2 expression in liver. **(a)** Graph indicating the Log2 fold depletion of the indicated EMC subunits in liver samples from LKO animals compared to WT controls. **(b)** Gene ontology on all proteins significantly downregulated in LKOs by a Log2 fold change of −1 or more using the Shiny Go algorithm^41^. **(c)** Volcano plot demonstrating the substantial reduction in FITM2 levels in EMC3 LKO animals. **(d**) WB corroborating the reduction in FITM2 protein expression in LKO liver lysates compared to WT samples (EGFR used as the loading control) **(e)** Densitometric quantitation of FITM2 depletion against the EGFR control. (**f)** HA-FITM2 stabilization with inhibition of proteasome (MG132, BTZ) or VCP/p97 (CB-5083) activity in U2OS FlpIn WT and ΔEMC6 cells. DOX-induced expression of HA-FITM2 over an 8 hr time course. **(g)** Steady-state levels of EMC subunits and clients (SQS, FITM2) in U2OS FlpIn WT, ΔEMC6 and ΔEMC6+rescue. Tubulin serves as a loading control. Each data point represents an individual animal. Statistical analysis: Shapiro-Wilk normal distribution, Outliers ROUT, and Unpaired Student t-test. The error bars represent the standard error of the mean (SEM).***p <0.001

### The EMC3-associated hepatic LD defect cannot be rescued by excess triglycerides

To further explore the hepatic fat storage-associated phenotype, we next fed the LKO mice *ad libitum* on a high fat diet for 9 weeks, i.e., under conditions of excess triglyceride availability. Interestingly, we found that the LKO animals were visibly leaner than littermate controls, despite having an equivalent food intake (**Fig. S2a-c**). Focussing on the mass of individual organs, we found that although the relative mass of the liver was not significantly different in LKO compared to WT mice, the weights of the two largest adipose tissue depots, inguinal (iWAT) and epididymal (eWAT) white adipose tissues were proportionately lower in the LKO animals compared to WT controls on HFD (**Fig. S2d**). This suggests, counter-intuitively, that hepatic deletion of EMC (which results in a reduced LD titre, **Fig.2h**) has a non-organ autonomous negative impact on the size of adipose tissue depots in the LKO. As on SD (**Fig. 2f**), we found that the HFD-exposed LKO mice had normal serum triglyceride levels but exhibited reduced fasting liver triglyceride levels (**Fig. S2e,f**). The liver of the LKO animals also appeared visibly less fatty relative to that of the WT control **Fig. S2g,h**). Taken together, our data indicate that LKO animals have a specific impairment in hepatic lipid storage, that cannot be rectified by increased triglyceride availability.

### FITM2, a key regulator of LD homeostasis, is depleted in EMC3-deficient livers

To gain insights into the candidate EMC client transmembrane proteins whose defective biogenesis may underpin the hepatic triglyceride storage/LD defects, we carried out a quantitative proteomic comparison of the proteins in WT and EMC3-deficient mouse liver samples (**Fig.3a-c**). As anticipated from previous studies^10^, levels of individual EMC subunits were substantially depleted (**Fig.3a**), while interrogation of the significant hits that were depleted by a Log2 fold change of −1 or more using the Shiny Go algorithm^41^ (**Fig.3b**) revealed an enrichment of proteins associated with various aspects of lipid homeostasis.

One of the most profoundly affected hits was the evolutionarily conserved protein FITM2 ^42^, whose depletion in *Emc3* KOs was confirmed by western blot of liver lysates (**Fig.3d,e**). FITM2 (also called FIT2, fat storage–inducing transmembrane protein 2) is an evolutionarily conserved polytopic ER-localized membrane protein that is required for LD homeostasis^42,43^. FITM2 depletion reduces LD accumulation in a range of contexts including cultured cells, adipocytes, Zebrafish, *C. elegans* and the fat body of *Drosophila melanogaster*^42,44–46^. By contrast, FITM2 overexpression promotes lipid droplet accumulation in a variety of models, including skeletal muscle^47^, mouse liver^42^ and in cultured insect cells^48^. We confirmed that FITM2 levels were reduced in a cell-autonomous manner using EMC6-deficient U2OS cells^9,10^ and that the FITM2 missing from EMC6 KO cells was being degraded by the proteasome (**Fig.3f**). Reintroducing EMC6 into EMC6 KO cells was sufficient to rescue endogenous FITM2 expression, reinforcing the requirement for an intact EMC for proper biogenesis (**Fig.3g**).

### FITM2 is the major EMC client in adipose tissues

FITM2 is expressed at high levels in mouse adipose tissues^42^, organs in which LDs play a key role, and is heavily implicated in lipid droplet homeostasis, fat storage and triglyceride binding^42–45,47^. Adipose tissues exist in two major forms: white adipose tissues (WAT), containing large unilocular LDs that act as an energy storage depot, and brown adipose tissues (BAT), a mitochondria-rich organ within numerous smaller lipid droplets, that catabolizes fatty acids, the product of lipolysis of LD-resident triglycerides to mediate non-shivering thermogenesis^31,49^. To test whether the EMC played a key role in adipose tissue homeostasis, we specifically deleted *Emc3* in adipocytes using adiponectin-cre^50^ to generate adipose tissue KOs (hence denoted as “AKO”). When we carried out an equivalent quantitative proteomic analysis on interscapular BAT (iBAT) and iWAT from these mice, once again we observed that along with the EMC subunits, FITM2 was substantially depleted (**Fig.4 a,b, Tables S5-7)**. We confirmed that FITM2 was depleted specifically in EMC3-deficient adipose tissues, but not in the liver samples isolated from the same animals (**Fig.4c**).

**Figure 4.**
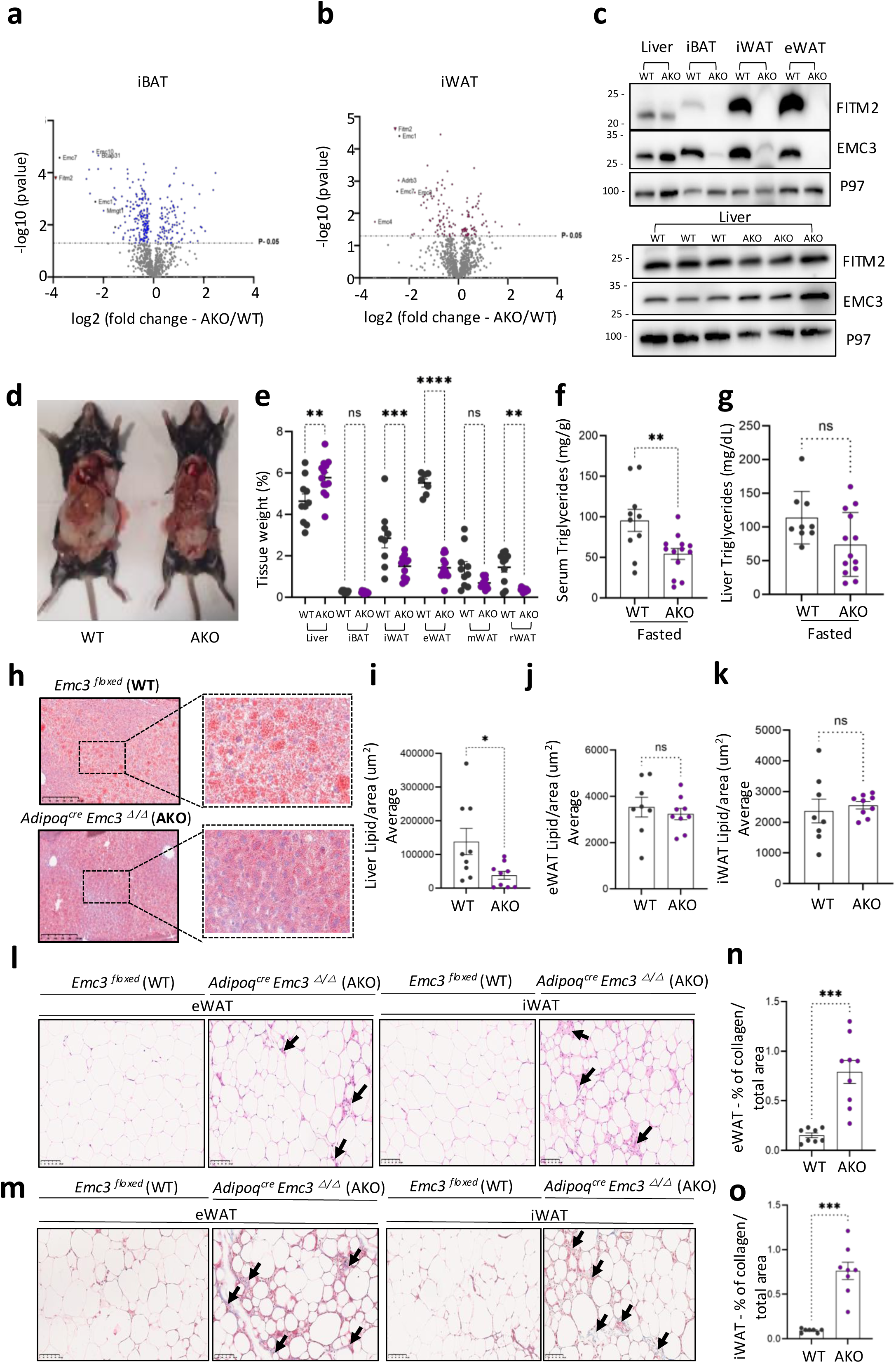
EMC3 adipose tissue-specific KOs (AKO) are viable but exhibit a reduced capability to store fat, reduced FITM2 expression and protection from obesity. Volcano plots demonstrating a substantial reduction in FITM2 protein levels in **(a)** intrascapular brown adipose tissue (iBAT) and **(b)** inguinal white adipose tissue samples from EMC3 AKO animals. **(c)** Representative WB panel comparing liver and adipose tissue from AKO animals versus WT controls. Depletion of EMC3 protein and reduced expression of FITM2 was confirmed by WB on samples from various adipose tissue depots (iBAT, iWAT, and eWAT). Liver samples from AKO animals were tested separately to confirm the adipose tissue specificity of the effect on EMC3 and FITM2 levels (p97 was used as a loading control). **(d)** Representative images from the abdominal cavity of WT and AKO animals exposed to HFD for 9 weeks from 3 weeks of age. AKO show reduced amount of eWAT compared to WT samples. **(e)** The same animals were evaluated for the weights of individual tissues. AKO animals present reduced accumulation of iWAT (inguinal white adipose tissue), eWAT (epididymal) and rWAT (retroperitoneal) weight and an increased liver weight compared to controls. **(f)** AKO animals present lower serum levels of triglycerides and **(g)** a tendency towards reduced levels of hepatic triglycerides. **(h)** Representative liver histopathology images (Oil red-O staining) and **(i)** statistical analyses corroborating the reduced accumulation of lipid droplets/area (10X magnification). Statistical analysis: Ordinary One-way ANOVA multiple comparisons/ Šidák’s multiple comparisons test. **(j-k)** No significant difference in lipid droplet size between the groups in the eWAT and iWAT. **(l)** Representative images of eWAT images and iWAT histopathology (H&E staining) from animals exposed to HFD for 9 weeks (20X magnification). Inflammatory cell infiltrates foci are indicated by the black arrows. The AKO animals exhibit higher inflammatory infiltration score, exhibiting a significant increase in the number of inflammatory cells within the eWAT and iWAT after 9 weeks on HFD. **(m)** Representative images (20X magnification) from eWAT and iWAT from animals exposed to HFD for 9 weeks stained with Masson’s Trichrome (MT) to reveal collagen deposits. The black arrows indicate increased collagen deposition compared to WT **(n,o)** Quantification of MT positive cells in eWAT or iWAT, corroborating the phenotype. Quantification of collagen deposition was realized using Qupath software using the plugin pixel classification by machine learning. Statistical analyses Outliers Rout, Shapiro-Wilk normality test - Unpaired Student t-test. Data from at least two biological replicates experiments each containing N=3 mice per genotype. The error bars represent the standard error of the mean (SEM). ns, non-significant (p>0.05);*p<0.05;**p<0.01;***p<0.001and ****p<0.0001.

### Deletion of EMC3 in adipose tissues drives lipodystrophy in response to obesogenic challenge

Focusing first on the response of the animals to standard chow conditions, as was reported for adipose tissue-specific deletion of *Fitm2*^51^, the body weights of young AKO adult mice were indistinguishable from control mice. While the relative liver mass of the AKO animals was significantly increased, the masses of individual adipose depots (**Fig. S3a,b)** and serum and hepatic cholesterol (**Fig. S3c,d**) were not significantly different than controls. While serum triglyceride levels were equivalent, the triglyceride levels found in livers of EMC3 AKOs were reduced (**Fig. S3e-f**). Hence, as encountered in the LKO animals, the deletion of EMC3 has non-autonomous impacts on other metabolic organs.

Miranda and colleagues reported that AT-specific FITM2 KO mice exhibited progressive lipodystrophy with ageing or in response to feeding with a high-fat diet (HFD)^51^. Therefore, we next challenged control versus EMC3 AKO mice with an experimental diet from which 60% of the energy is derived from fat^52^. Following a short period (9 weeks) of *ad libitum* feeding on HFD, the body cavities of the EMC3 AKO mice exhibited less adiposity than controls (**Fig.4d**), with the under-representation of multiple adipose tissue depots in AKO animals (**Fig.4e**). Similar phenotypes have been reported for FITM2 adipose tissue-specific KOs^51^. This HFD-induced lipodystrophy could not be explained by differences in food intake (**Fig.S3g**). Strikingly, AKO mice had reduced levels of serum triglycerides (**Fig.4f**). As observed in a more pronounced manner in the animals on standard chow (**Fig.S3f**) AKOs also, somewhat counterintuitively, showed a tendency towards reduced hepatic accumulation of triglycerides (**Fig.4g-i**) albeit with a significantly increased liver mass, as observed in FITM2 AKOs^51^.

Although depletion of *Fitm2* impacts LD size or titre in several models^43,47,48,51,51,54^, the primary impact on WAT is a pronounced reduction in adipocyte numbers and hence adipose tissue mass^51^. Strikingly, although loss of FITM2 does not impinge substantively on LD size of the remaining adipocytes, the remaining WAT exhibits extensive inflammatory infiltrates, likely associated with adipocyte death^51^.

Similarly, while the size of the inguinal and epididymal white adipocytes from the surviving WAT in EMC3 AKOs was not appreciably different in size from controls (**Fig4.j,k**), there was however evidence for extensive immune infiltrates in WAT from EMC3 AKOs (**Fig.4l**) and collagen deposition (**Fig.4m**) that is indicative of fibrosis (**Fig.4 n,o**). Hence, like the FITM2 KO models^51^, the deletion of EMC3 impacts on adipose tissue homeostasis by causing adipose tissue lipodystrophy and adipose tissue inflammation, in response to obesogenic challenge.

### Prolonged feeding on HFD causes profound lipodystrophy in older EMC3 AKO animals

To further study the impact of ageing on EMC3 AKO animals, we next examined the effect of prolonged feeding on normal chow or HFD for 29 weeks, at which point the animals had aged to ∼8 months. Notably, as observed in FITM2 KO animals^51^, while the overall body weight of standard chow-fed EMC3 AKO animals was unaffected (**Fig.5a**), the body cavity of these animals appeared leaner than controls (**Fig.5b**). Indeed, the visceral (epididymal) adipose depot mass was significantly reduced, while the other adipose depots were normal (**Fig. 5c**). This mild lipodystrophic phenotype was dramatically exaggerated in animals exposed to HFD. AKO animals, which largely resisted HFD-induced weight gain (**Fig.5a**) showed a remarkably reduced body cavity adiposity (**Fig.5d**) which was associated with significant reduction to overall fat mass, particularly in the iWAT and eWAT depots (**Fig 5e-g**). The remaining WAT that survived in the AKO mice on HFD exhibited a significant reduction in LD size (**Fig.5h-i**) and significant deposition of collagen (**Fig.5j**).

**Figure 5.**
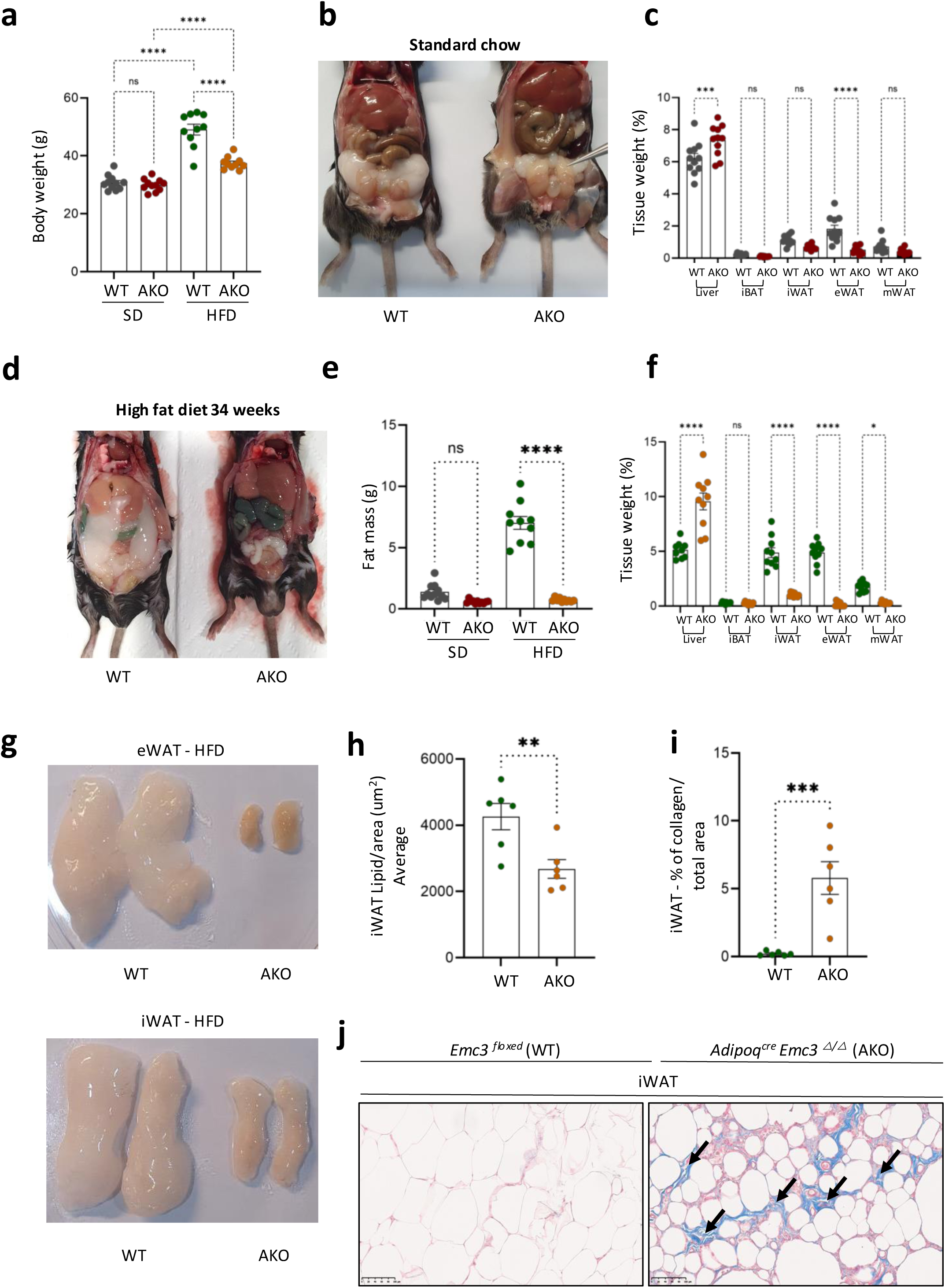
Under prolonged HFD feeding, AKO animals show pronounced lipodystrophy and collagen deposition, reduced body weight and fat mass. **(a)** AKO animals fed for 28 weeks *ad libitum* on HFD exhibit lower body weight compared with the WT controls **(b)** Although AKO animals fed *ad libitum* for the same time on SD do not exhibit differences in body weight, they exhibit reduced accumulation of fat in their abdominal cavity and **(c)** demonstrate significant reduction in eWAT mass and increased liver weight **(d).** Representative images from the abdominal cavity of WT and AKO animals exposed to HFD for 28 weeks. AKO animals show **(e)** a significant decrease in fat mass and **(f-g),** a pronounced reduction in adipose tissue accumulation in iWAT, eWAT and a higher liver mass. Statistical analyses: Ordinary One-way ANOVA multiple comparisons/ Tukey’s multiple comparisons test. **(h,i)** Reduced lipid droplet size compared to WTs and (j) iWAT from animals exposed to HFD present elevated collagen deposition,(MT positive staining). Collagen spots indicated by the black arrows. Quantification of collagen deposition was realized using Qupath software using the plugin pixel classification by machine learning. Statistical analyses Outliers Rout, Shapiro-Wilk normality test - Unpaired Student t-test. Data from at least two experiments each containing N=3 mice per genotype. The error bars represent the standard error of the mean (SEM). ns, non-significant (p>0.05);*p<0.05; **p<0.01; *** p<0.001; ****p<0.0001.

EMC3 AKO mice also exhibited a substantially increased liver mass both on standard chow and HFD (**Fig.5c,f**). Although these differences could not obviously be attributed to increased fat deposition in AKO mice livers (**Fig.S4a-d**), these livers exhibited evidence of elevated collagen deposition that was similar to their adipose depots (**Fig.S4e-g**). These pronounced phenotypic differences could not be attributed to significant differences in food intake (**Fig.S4g**). We conclude that akin to deletion of FITM2^51^, loss of EMC3 in adipose tissues results in profound lipodystrophy, fat storage and partitioning defects, fibrosis and inflammation.

### Adipose tissue-specific deletion of EMC3 results in hypertrophic LDs in BAT

We next focused on the impact of EMC3 depletion on appearance of the BAT - an organ that plays a key role in whole-body metabolism and specifically, non-shivering thermogenesis^49^. In stark contrast to WAT, loss of EMC3 did not appreciably impact the mass of intrascapular BAT in animals on standard chow (**Fig. S4b**) or on HFD (**Fig.4e, Fig.5c)**. However, we observed a marked enlargement in the size of individual LDs within the brown adipocytes (**Fig. 6a,b**). This LD hypertrophic phenotype of the EMC3-deficient BAT compared to WT controls was sustained when mice were maintained at 30°C (i.e., thermoneutrality) or 4°C (i.e. conditions that promote non-shivering thermogenesis), suggesting that it may have an impact on normal BAT physiology. This enlarged LD phenotype in BAT has been reported for mice deficient in FITM2^51^, whose levels were dramatically depleted in EMC3-deficient iBAT (**Fig.4a,c**).

**Figure 6.**
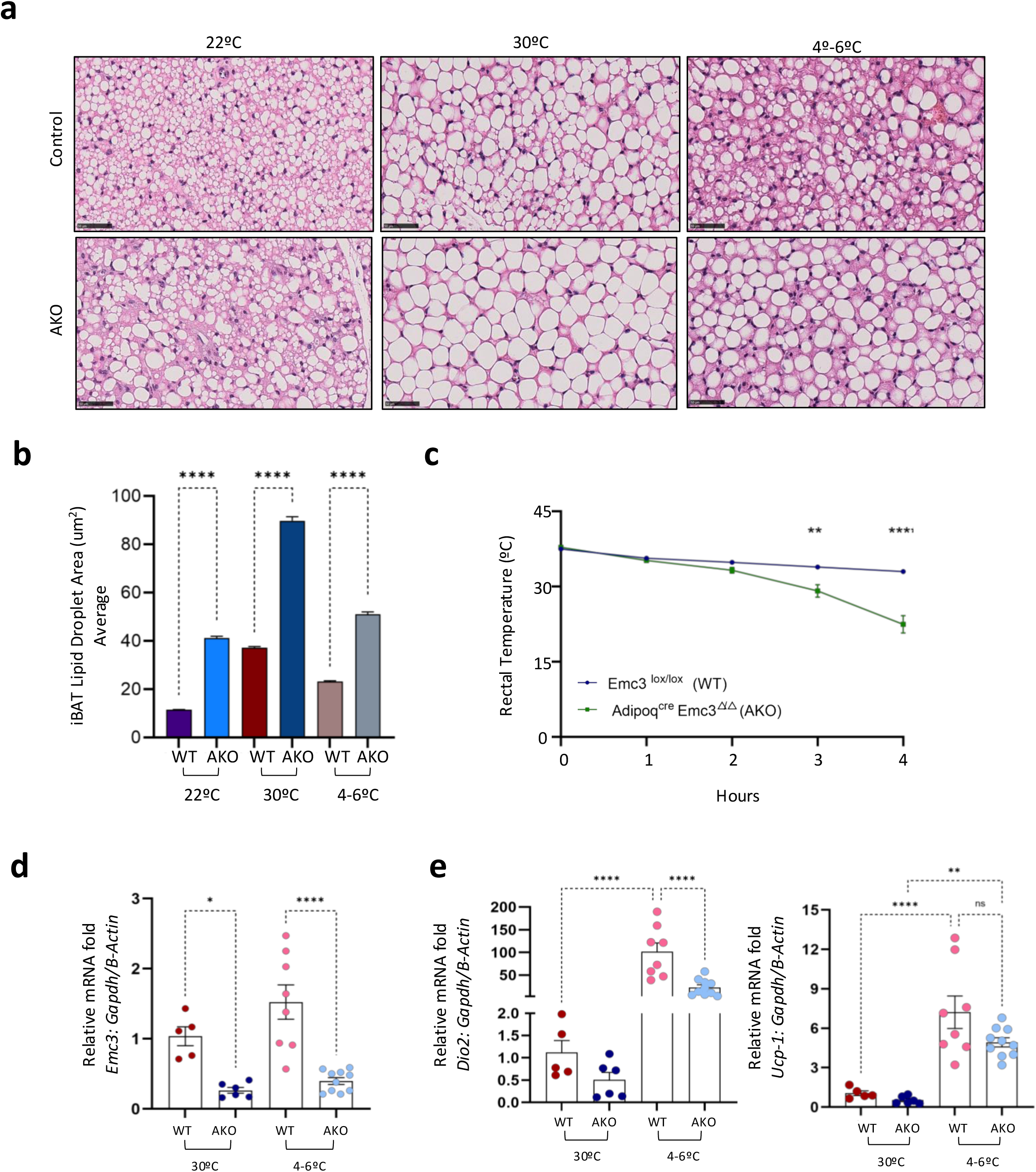
AKO animals exhibit hypertrophic lipid droplets in iBAT, high cold sensitivity and perturbations in expression of thermogenic genes during cold stress. **(a)** Representative H&E-stained images from iBAT demonstrating lipid droplet hypertrophy in AKO animals (magnification 40X) compared to WT controls under all temperature regimes. **(b)** Significant differences in LD area between WT and AKO mice corroborates the AKO-associated lipid droplet hypertrophic phenotype. Statistical analysis: Ordinary One-way ANOVA multiple comparisons/Tukey’s multiple comparisons test. **(c)** Under acute cold exposure (4-6°C), AKOs present significantly reduced rectal temperature after 3 and 4 hours of cold exposure. The individual variance for each timepoint was taken into consideration for this analysis. Statistical analysis: Multiple Unpaired Student t-test analyze. Method Two-Stage step-up (Benjamini, Krieger, and Yekutieli). Data from at least two biological replicates experiments each containing N=3 mice per genotype. The error bars represent the standard error of the mean (SEM). **p<0.01; ****p<0.0001**. The** mRNA levels confirm the downregulation of (d) *Emc3* mRNA in iBAT from AKO animals under both thermoneutrality and cold. Under cold conditions (4°-6°C), AKOs present a significant decrease in mRNA levels of (e) *Dio2* and elevated expression and no significant difference in *Ucp-1* mRNA levels when compared with WT controls under thermoneutrality or cold exposure. Cold exposure data are from three biological replicate experiments each containing N=3 mice per genotype while the experiments done under thermoneutrality are from two biological replicate experiments each containing N=3 mice per genotype. Statistical analyses: Ordinary One-way ANOVA multiple comparisons/ Tukey’s multiple comparisons test. The error bars represent the standard error of the mean (SEM). ns, non-significant (p>0.05); *p<0.05; **p<0.01; and ****p<0.0001.

### AKO mice exhibit cold sensitivity

Thermogenic signals, including cold, activate lipolysis in adipose tissues: the breakdown of triglycerides within LD into glycerol and fatty acids^49^. In the BAT, free fatty acids activate UCP1 (uncoupling protein-1), a protein that spans the inner mitochondrial membrane. UCP1 uncouples oxidative phosphorylation from ATP synthesis by facilitating proton leakage back across the mitochondrial inner membrane; this futile biochemical cycle of ATP dissipation generates heat^49^. To ascertain the thermogenic capacity of the EMC3 AKO mice, we preconditioned the mice at thermoneutrality (30°C, i.e., under conditions where there is no thermogenic demand), then exposed then to cold (4-6°C). Strikingly, AKO mice were profoundly defective in their ability to sustain core body temperature in response to cold challenge (**Fig. 6c**). This is consistent with the AKO-associated LD and adipose tissue perturbations (**Figs. 4-6**), particularly in WAT, which plays a critical role in the provision of fatty acids for BAT thermogenesis^55^. Interestingly, we observed aberrant expression the thermogenic gene Dio2, in EMC3 KO BAT (**Fig. 6d-e**) whereas the levels of UCP1 mRNA was not significantly altered (**Fig. 6e**). Our data show that the EMC plays a critical role in coordinating white adipose tissue homeostasis and is required to support thermogenesis.

### Ablation of the *Drosophila* EMC3 homolog dPob results in defects in LD homeostasis

As the EMC is conserved in eukaryotes^42^, we hypothesised that its role in LD homeostasis, revealed by our mouse studies, may be conserved in other metazoans. To test this, we selected the fat body of *Drosophila melanogaster*, a LD-rich key fat storage organ that fulfils analogous roles to both the mammalian liver and adipose tissues.

Clones of fat body cells homozygous for a deletion in the *Drosophila* EMC3 ortholog dPob, labelled by the absence of nuclear GFP expression, exhibited pronounced defects in LDs including a dramatic reduction of the Nile red-reactive signal (**Fig.7a,b**) and a significant reduction to LD area and perimeter (**Fig.7c,d**). Moreover, the remaining LDs in dPob/EMC3-deficient territories were smaller and more numerous than in the adjacent WT tissue (**Fig.7a,b,e**). This appeared similar to the phenotype observed upon ablation of *Drosophila* FIT2 in the fat body^46^. We next examined whether mouse FITM2 was recognised as an EMC client in *Drosophila* cells, focusing on a comparison between transgene-driven HA-FITM2 expression in WT versus adjacent dPob/EMC3-deficient clones in the *Drosophila* imaginal eye disc. Expression of FITM2-HA was substantially reduced in dPob/EMC3-deficient mitotic clones compared to adjacent WT GFP-positive territories (**Fig.7f**), indicating an intrinsic EMC-dependence for biogenesis.

We next examined the molecular determinant(s) within mouse FITM2 that conferred its EMC-dependent insertion. Inspection of the six TM helices of mouse FITM2 (**Fig.S5a**) revealed that its first TMD contained several atypical residues that resulted in a reduced overall score on the Zhao-London hydrophobicity scale^56^. Reduced hydrophobicity has been identified as one factor contributing to EMC dependency of some tail-anchored proteins^9,10^ which have a similar cytoplasmic N-terminal topology to the first TMD of FITM2. To investigate this, we mutated three non-hydrophobic residues (His 23, Glu 40, Ser 42), plus the proline residue in this TMD (Pro 25), to the more hydrophobic leucine (L) residues (**Fig.S5b**). While mutating two non-hydrophobic residues (Glu 40, Ser 42; FITM2-2L) positioned in the luminal-facing third of the TMD had little impact, a mutant in which all four residues were replaced by leucines (FITM2-4L) was readily expressed in EMC3 mutant clones, effectively circumventing EMC-dependent biogenesis (Fig. 7f-g). Expression of another EMC client, the Na^+^K^+^ ATPase^11,14^ remained compromised in EMC3 mutant clones **Fig S5c**). This suggests that low hydrophobicity of FITM2’s first TMD is a determinant that renders it dependent on the EMC for its biogenesis.

**Figure 7.**
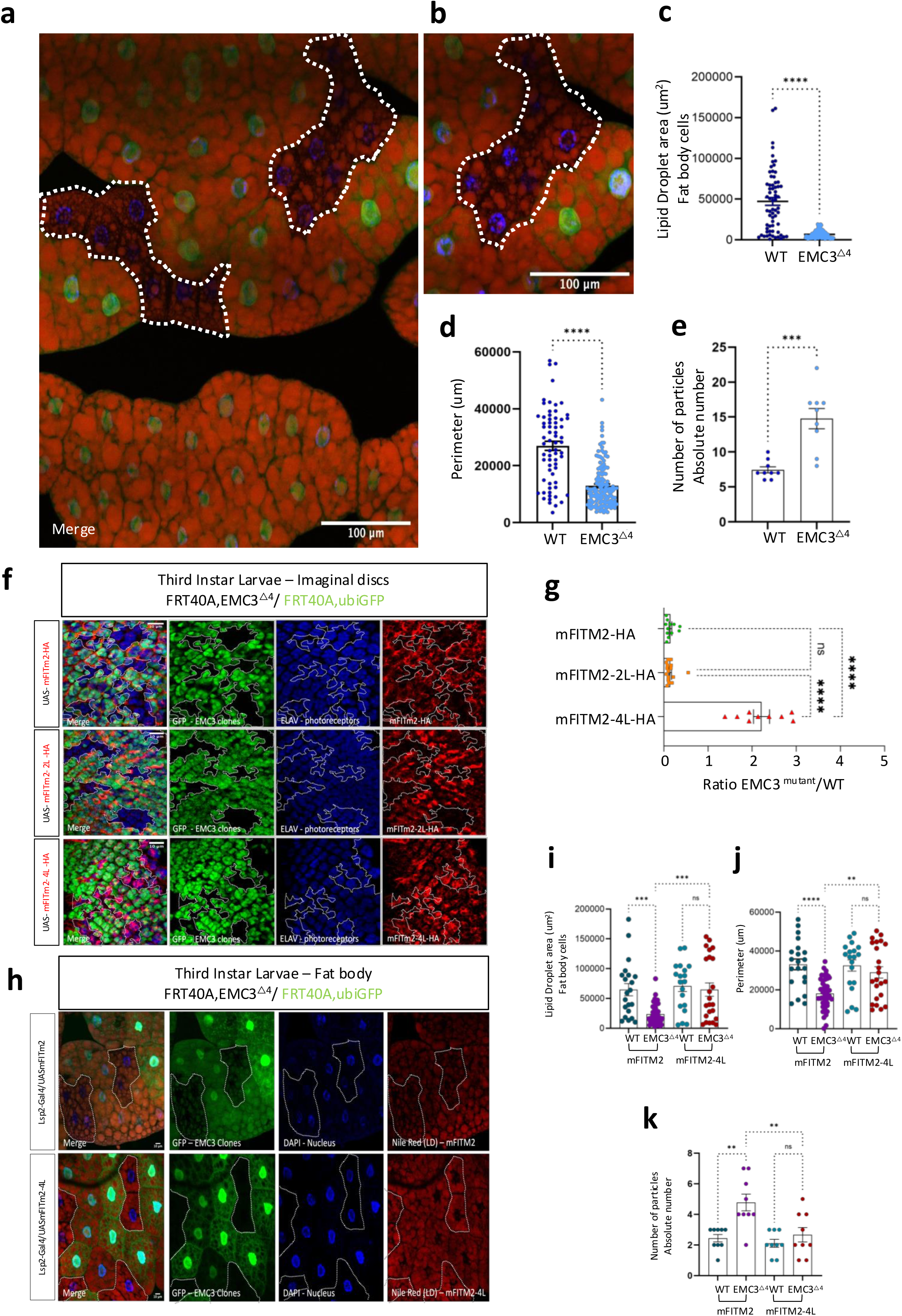
Ablation of EMC3 in *Drosophila* fat body cells causes a lipid droplet homeostasis defect that can be rescued with FITM2-4L, an EMC-independent mutant form of mFITM2. **(a,b)** Representative confocal images of the fat body cells from third instar larvae of *Drosophila melanogaster* mosaic tissue. EMC3^△4^ mutant clones are labeled by the absence of GFP expression (delimited by the white dotted lines). DAPI was used to stain the nucleus and Nile Red to stain lipid droplets. A significant difference is observed in the lipid droplet size and number between the EMC3^△4^ mutant cells, exhibiting reduced **(c)** lipid droplet area and **(d)** reduced perimeter compared to WT cells. Furthermore, EMC3^△4^ mutant cells show **(e)** an increase in individual Nile Red-positive particles than WT cells. Lipid droplet quantification was determined using, at least, three patches of WT and mutant cells per fat body image, using ImageJ. Three images from different flies were used (A total of 9 images analyzed per genotype). Statistical analyses: Outliers ROUT; Normality and Lognormality tests - Shapiro-Wilk normality test. T-tests (and nonparametric tests) - Unpaired T-test / Mann-Whitney test. Error bars correspond to the standard deviation (stdev) of the mean. **(f)** Representative confocal images of eye imaginal discs from third instar larvae of *Drosophila melanogaster* mosaic tissue containing EMC3^△4^ mutant clones labeled by the absence of GFP expression (delimitated by the white dotted line). The expression of mFITM2, mFITm2-2L, and mFITm2-4L is revealed by anti-HA staining (red) which anti-ELAV staining was used to label the photoreceptors (blue). The upper panel shows the reduced expression of WT mFITM2-HA variant in EMC3^△4^ homozygous territories. The middle panel shows reduced expression of mFITM2-2L-HA in EMC3^△4^ homozygous territories. The bottom panel bottom shows that territories expressing mFITM2-4L-HA exhibit an increased immunofluorescent intensity compared to WT or mFITM2-2L-HA-expressing cells. **(g)** Quantification of the ratio of fluorescence intensity for the HA (red) channel between EMC3^△4^ homozygous mutant territories and neighboring non-mutant cells for mFITM2, mFITM2-2L, and mFITM2-4L. A statistically significant difference was observed for mFITM2-4L compared to mFITM2 or mFITM2-2L. At least three patches of WT and mutant cells were quantified per eye imaginal disc. Three eye imaginal discs from different flies were used (N=3). Each data point represents an individual image analyzed. Statistical analysis: Ordinary One-way ANOVA - Dunn-*Šidák* multiple comparison test. Error bars correspond to the standard deviation (stdev) of the mean. **(h)** Representative confocal image of the fat body from third instar larvae of *Drosophila melanogaster* stained with Nile-red. The upper panel shows mosaic tissues containing EMC3^△4^ mutant clones (labeled by the absence of GFP expression) expressing WT mFITM2-HA or expressing the mFITM2-4L-HA mutant (bottom panel). Note that the territories expressing the 4L variant demonstrate a rescue of the EMC3^△4^ mutant phenotype**. (i-k)** Lipid droplet quantification was determined using, at least, three patches of WT and mutant cells per fat body image, using ImageJ. Three images from different flies were used (A total of 9 images analyzed per genotype). Statistical analyses: Outliers ROUT; Normality and Lognormality tests - Shapiro-Wilk normality test. T-tests - Unpaired T-test. The error bars represent the standard error of the mean (SEM). ns, non-significant (p>0.05); **p<0.01; ***p=0.001 and ****p <0,0001.

Since EMC3 ablation profoundly affects LD homeostasis in the fat body (**Fig.7a-e**), we next examined the impact of overexpression of WT FITM2 versus FITM2-4L on this EMC3-associated defective LD phenotype. Expression of WT FITM2, whose biogenesis is not supported in EMC3-deficient territories, had no impact on the mutant LD phenotype (**Fig. 7h**). In contrast, the FITM2-4L was able to rescue the LD defect (**Fig.7h-l**). Taken together, our data establish that the EMC is required for LD homeostasis and that one of its key client proteins, FITM2 requires the EMC to support the biogenesis of non-hydrophobic features contained within its first TMD.

## Discussion

Bona fide client proteins of the EMC exhibit a spectrum of dependency on it for insertion and biogenesis. Therefore, ablation of the EMC in cells/tissues does not necessarily stoichiometrically deplete its client proteins and can result in residual expression [refs]. Consequently, it can be challenging to relate whether the extent of depletion of an EMC client protein impacts a biological process substantially in a given tissue. To establish which biological processes are strongly dependent on membrane protein insertion by the EMC, we undertook an organismal approach that coupled tissue specific phenotypes with a proteomic screen to identify essential EMC client proteins, and focusing on both liver and adipose tissues. These complementary objective approaches allowed us to systematically identify a critical role for the EMC in LD homeostasis through governance over insertion of FITM2. FITM2 emerged as the most profoundly EMC-dependent integral membrane proteins in the liver, iBAT and iWAT, whose depletion was comparable or greater than the EMC subunits themselves (**Figs. 3,4, Tables S5-7**). Hence, our approach successfully identified a highly sensitive client protein of the EMC and a cohort of phenotypes that resemble those associated with loss of this novel client protein, FITM2. By demonstrating that the EMC is required for LD homeostasis in the *Drosophila* fat body - an organ that serves analogous roles in lipid homeostasis to the mammalian liver and adipose tissues^57^, we show that the EMC‘s role in metabolic regulation is evolutionarily conserved.

Although the EMC fulfils pleiotropic biological roles, and while we do not exclude a role for other potential clients, our data reveals a striking similarity between the EMC3-associated phenotypes observed in our study and the previously identified phenotypes of mice deficient in FITM2, a key regulator of LD homeostasis^42–45,47^. Several of the major phenotypes observed in our EMC3 AKO resemble those found in the equivalent FITM2 KO studies (e.g. lipodystrophy, enlarged LD in the BAT, impaired triglyceride homeostasis).

There are also similarities between phenotypes observed in our LKO model and those reported for FITM2 LKO animals^58,59^. These include hepatocyte cell death, liver function and derangements in lipid homeostasis (**Figs. 1,2 S1,2**)^59^. There appears to be a greater congruence in the phenotypes associated with exposure of these animal models to high fat diet^59^. As with FITM2 LKO animals, our model displayed reduced hepatic triglycerides, a trend towards reduced hepatic Oil Red O staining, and reduced body weight of the LKOs compared to controls ^59^.

Intriguingly, we also observe that hepatic depletion of EMC3 results in an overall reduced body weight on HFD, with reduced adiposity in the gonadal and inguinal adipose tissue depots, as Bond and colleagues did for FITM2 LKOS. Like FITM2, EMC3 (and by extension the EMC) plays a role in driving non-organ autonomous impact on fat storage in adipose tissues. However, our LD-associated phenotypes observed in the LKO model differ substantially from the reported liver-specific FITM2 KO phenotypes, particularly on standard chow^58,59^. This is reflected by increased hepatic triglycerides and cholesterol^58,59^, while our study observed reduced triglyceride levels in liver.

One explanation for some of these apparent discrepancies is that while FITM2 is the most profoundly affected client protein in our model (**Fig.3c, Fig.4a,b**), additional EMC clients may contribute to the death of EMC3-deficient hepatocytes and/or some of the contradictions observed between our LKO animals and the FITM2 LKOs on SD. We believe that the relatively mild–and notably transient–phenotypes observed in the EMC3 LKO animals are a result of death of EMC3-deficient hepatocytes, followed by their replacement with WT hepatocytes that arise and proliferate following incomplete Cre excision, as has been reported in other studies^60^.

It is interesting that the loss of EMC3, like the depletion of FITM2, has its most profound impact on WAT - driving pronounced lipodystrophy. In contrast, the BAT in the EMC3 and FITM2 AKO animals exhibit hypertrophic LDs, without causing atrophy. Although it is difficult to separate cell-autonomous, versus indirect impacts in an organismal model, we favour the hypothesis that the loss of white adipocytes places pressure on the adipocytes in BAT to act abnormally as a secondary triglyceride store, driving LD hypertrophy and whitening as a side effect. This is predicted to have a critical impact on non-shivering thermogenesis. The reduction in WAT, normally the major source of fatty acids will place more pressure on the ability of the BAT to generate fatty acids via lipolysis to support UCP1-driven thermogenesis. However, having undergone whitening, the BAT of EMC3 AKOs may be less competent to support non-shivering thermogenesis. This is evidenced by the reduced capacity to sustain core body temperature and the perturbed expression of several thermogenesis-associated genes in the AKO BAT (**Fig.6**). Interestingly, the ablation of EMC10, which exists in two isoforms: an integral membrane protein that forms part of the EMC, versus a smaller secreted form, results in similar metabolic defects, including reduced adiposity and resistance to high fat diet^61^.

While our data strongly favour the hypothesis that the LD defects observed upon loss of the EMC are driven by loss of FITM2, another possibility is that EMC depletion impacts the biogenesis of LD-resident proteins that impinge on LD homeostasis. These proteins contain hydrophobic segments or hairpin-like structures that mediate membrane association. The EMC-dependency for a small subset of LD proteins has been proposed from *in vitro* studies^62^, although it remains unclear how profoundly affected these are *in vivo*, and whether this is sufficient to drive the LD-associated phenotypes observed here. Although we cannot fully rule out this possibility, our proteomic data show that LD-associated proteins identified by Leznicki and colleagues, as well as key LD homeostasis regulators like perilipins, are not significantly affected in our EMC3 KO model, in contrast to the profound reduction in FITM2 levels **(Table S8)**.

While we observed tissue-autonomous LD phenotypes in the LKO and AKO models, both models showed that EMC3 depletion affected metabolic crosstalk between the liver and adipose tissues and vice versa. On one level, this is to be expected, since anything that impinges on fat storage and triglyceride homeostasis in one organ will impinge upon lipid homeostasis in others. Future studies will be required to establish the basis of this defect. The precise mechanistic role(s) of FITM2 that explain its profound impact on LD homeostasis also remain(s) to be fully delineated. The proposed roles range from recruiting proteins associated with the cell biology of LD formation^63^, or triglyceride binding^48^ or in controlling the directionality of LD emergence towards the cytoplasm^58^. However, a critical aspect that requires further attention is that FITM2 and its orthologs harbour a lipid phosphatase enzymatic activity in the ER lumen that is critical for some of FITM2’s biological functions, including LD formation^44,64^. This fatty acyl-coenzyme A (CoA) diphosphatase activity has been proposed to mediate the hydrolysis of unsaturated fatty acyl–CoA species and may explain why the latter are normally asymmetrically found on the cytoplasmic face of the ER ^44^. It remains to be determined what biological roles either FITM2-associated substrate(s) or products fulfil, and how these influence LD homeostasis in a cell-autonomous manner and/or metabolic organ cross-talk.

To probe the potential conservation of EMC functions amongst metazoans, we exploited a *Drosophila* model to map the determinants of murine FITM2 that confer EMC-dependency. Although we note that several other features beyond hydrophobicity pertain to polytopic membrane protein EMC clients^8^, we were able to show that a more hydrophobic version of the first TMD of FITM2 TMD rendered murine FITM2 EMC-dependent **(Fig 7)**. This allowed the FITM2-4L mutant to be expressed in EMC3-deficient cells in the *Drosophila* imaginal disc and in the fat body, where FITM2-4L rescued the EMC3-associated LD defect in that tissue. Overall, our data identify a critical and conserved role of the EMC in LD homeostasis though the membrane insertion of FITM2.

**Supplemental Figure 1.**
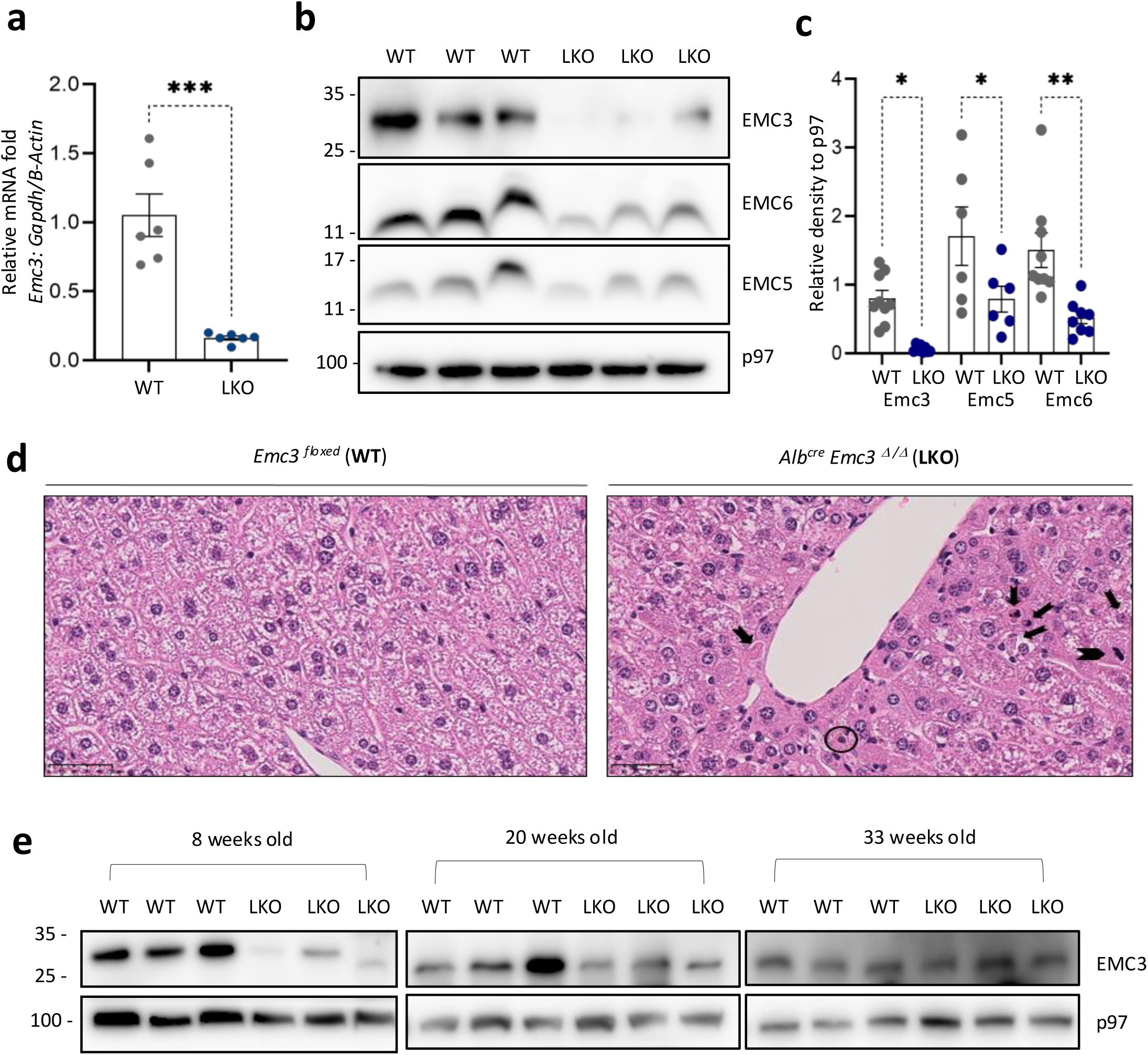
EMC3 KO liver-specific deletion affects the expression of other EMC core subunits and causes hepatocyte cell death and compensatory proliferation. **(a)** LKOs exhibit reduced hepatic *Emc3* mRNA expression compared to the WT group. *Gapdh* and *b-Actin* are used as reference genes. Statistical analyses: Shapiro-Wilk normality test - Unpaired Student t-test. **(b)** LKO animals exhibit reduced EMC6 and EMC5 protein expression. A p97 immunoblot is the loading control). **(c)** Densitometric quantification of WB images by Image J, based on an ROI selection; p97 expression was used as the normalization factor. Statistical analysis: Ordinary One-way ANOVA multiple comparisons/ Šidák’s multiple comparisons test. **(d)** Liver samples were stained with H&E. LKO liver samples show evidence of cell death (necrosis/apoptosis), indicated by the black arrows. The black circle shows ceroid pigment (a homogenous smudge in pink), indicating macrophage activity. The arrowhead indicates mitotic cells. Images were captured at 40X magnification. **(e)** Western blotting indicates an increase in EMC3 expression in liver samples of LKO animals at 20 and 33 weeks, old compared to 8-week-old animals. Statistical analyses: Shapiro-Wilk normality test Outliers ROUT and Unpaired Student t-test. Data are obtained are from, at least, two distinct biological replicate experiments each containing N=3 mice per genotype. Each data point represents an individual animal. The error bars represent the standard error of the mean (SEM). ns, non-significant (p>0.05); *p<0.05; **p<0.01 and ***p<0.001.

**Supplemental Figure 2.**
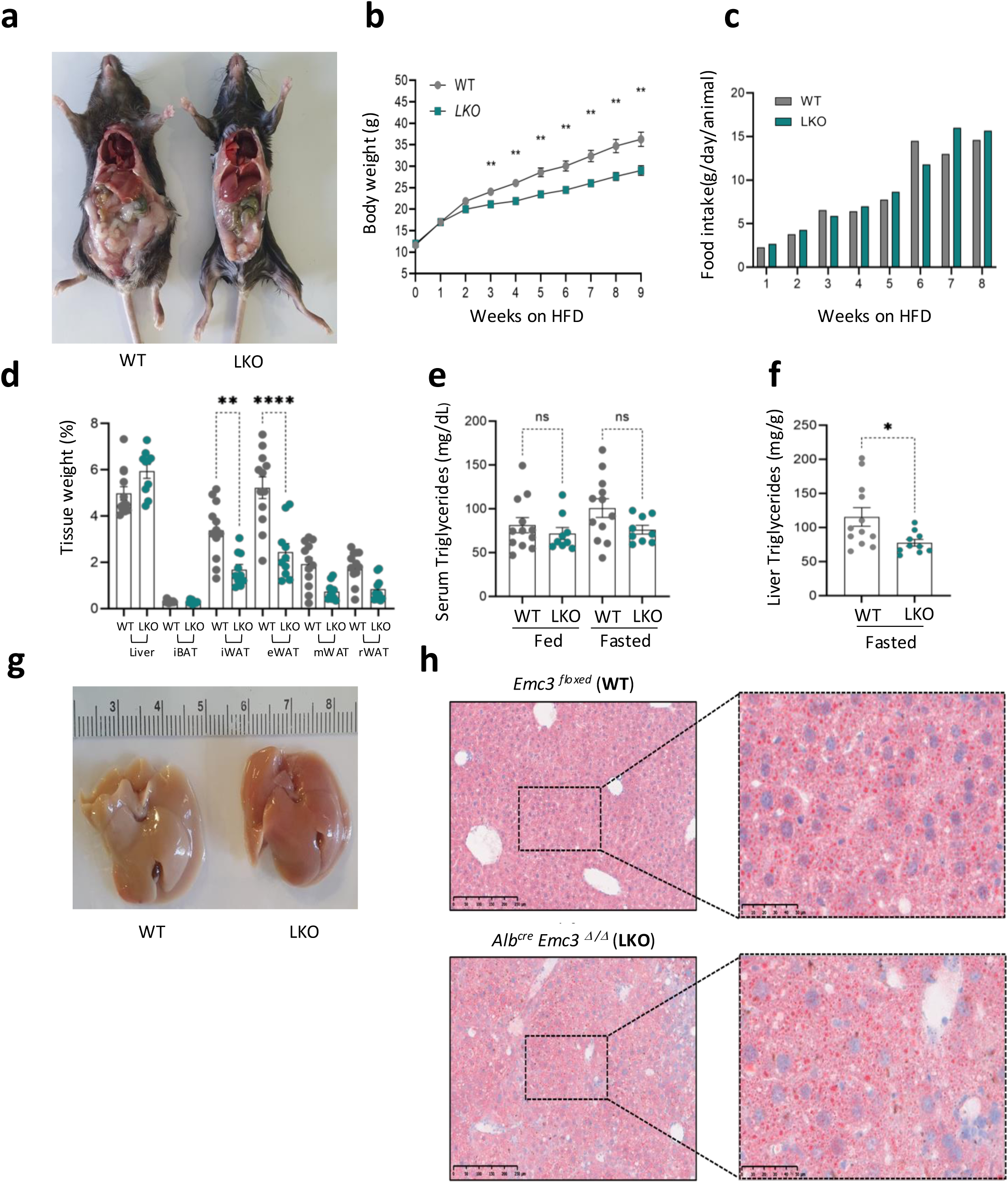
Metabolic effects in LKO animals after HFD challenge. **(a)** LKO animals on 9 weeks *ad libitum* on HFD exhibited reduced accumulation of fat in their abdominal cavity compared to WT animals. **(b)** Marked difference in LKO body weight measurement weekly through 9 weeks on High Fat Diet (HFD) exposition. Statistical analysis: multiple Unpaired Student t-test analyze. Method Two-Stage step-up ^65^ **(c)** No significant difference in food intake between the groups. **(d)** LKOs show significant reduction in iWAT (inguinal white adipose tissue) and eWAT (epididymal) weights. **(e)** LKO animals show no significant difference in serum triglycerides levels. Statistical analysis: ordinary one-way ANOVA multiple comparisons/Tukey’s multiple comparisons test. **(f)** Reduced levels of hepatic triglycerides in LKO animals after fasting. Statistical analyses involved normality and lognormality tests followed by an Unpaired Student t-test. **(g-h)** Representative photographs of livers and representative Oil Red-O liver-stained images from WT and LKO animals. Data from at least two experiments, each containing N=3 mice per genotype. The error bars represent the standard error of the mean (SEM). ns, non-significant (p>0.05); *p<0.05; **p<0.01; ****p<0.0001.

**Supplemental Figure 3.**
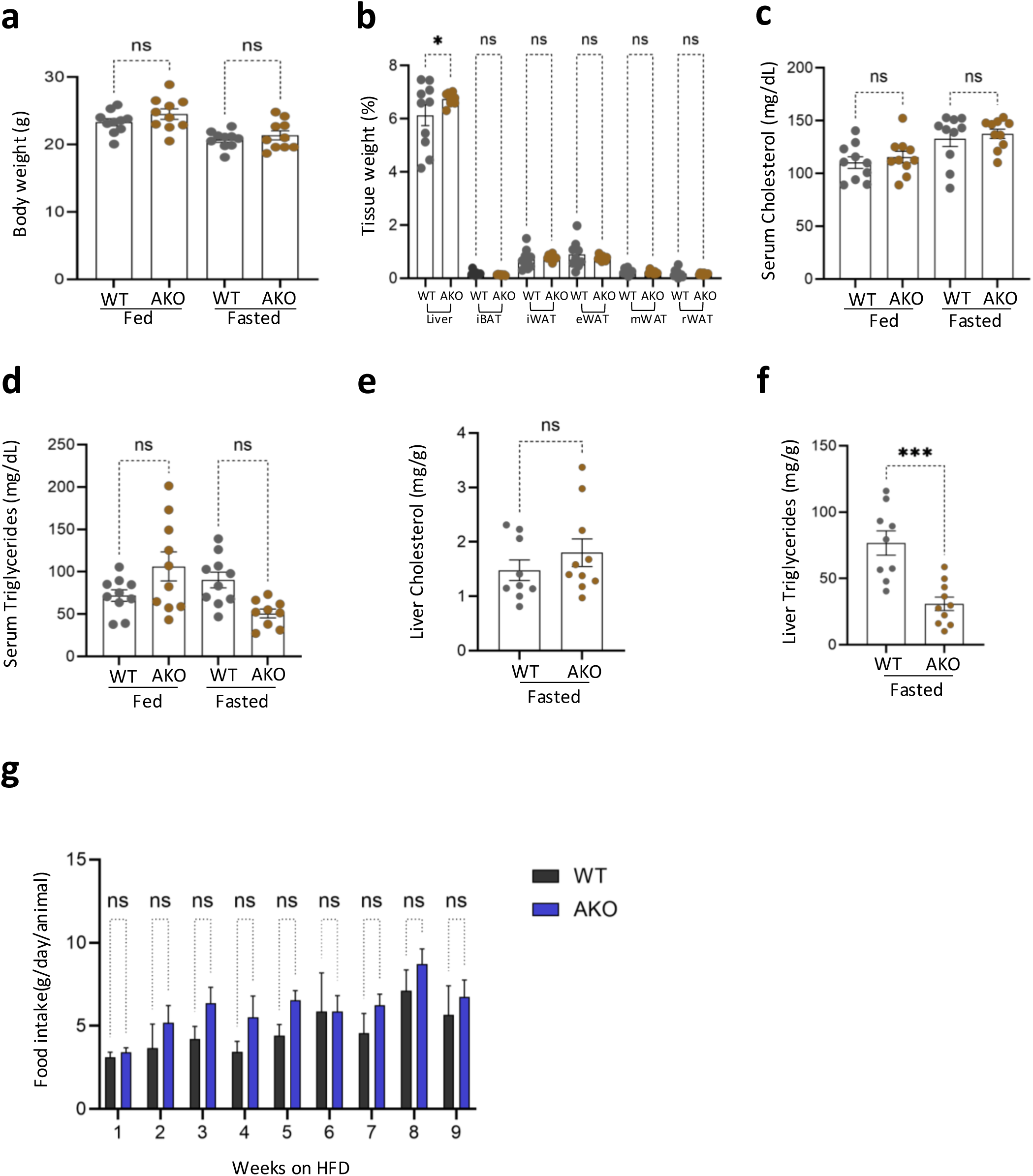
AKO animals fed with SD for 9 weeks exhibit significant difference in liver triglycerides compared to WT animals. AKO maintained for 9 weeks *ad libitum* on SD do not present a significant difference in **(a)** body weight, **(b)** adipose tissue weight**, (c)** serum cholesterol, or **(d)** serum triglycerides. Statistical analyses: Ordinary One-way ANOVA multiple comparisons/ Šidák’s multiple comparisons test. **(e)** AKO livers do not present a significant difference in liver cholesterol but **(e)** exhibit significantly reduced triglyceride levels after fasting. (g) No significant difference in food intake between the groups. Statistical analyses: Outliers Rout, Shapiro-Wilk normality test - Unpaired Student t-test. Data from three experiments, each containing at least N=3 mice per genotype. The error bars represent the standard error of the mean (SEM). ns, non-significant (p>0.05); *p<0.05; and ***p<0.001.

**Supplemental Figure 4.**
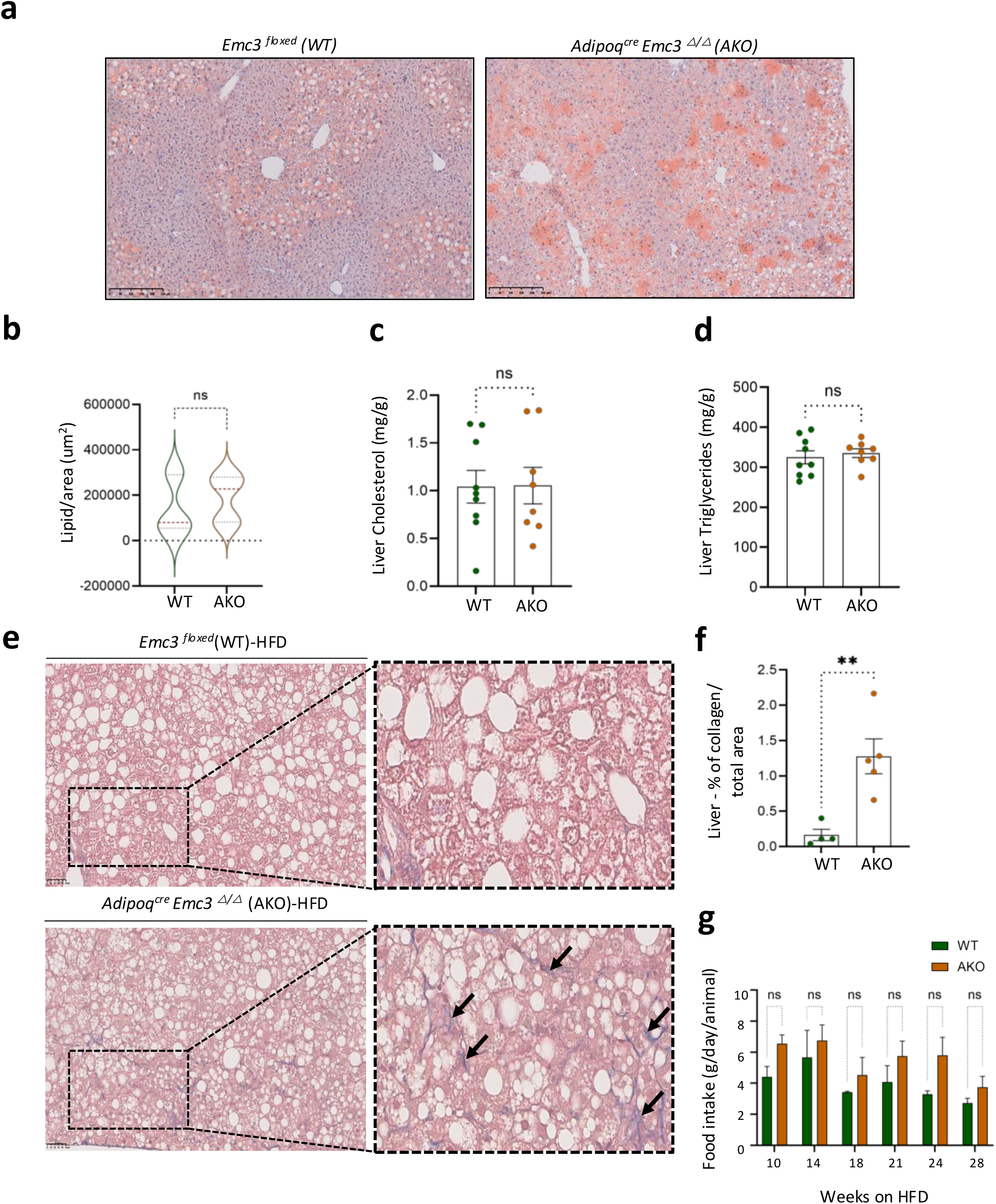
AKO animals show hepatic collagen accumulation under long term HFD exposure. **(a)** Representative liver histopathology images (Oil red-O staining) (20X magnification), to detect hepatic lipid deposition. **(b)** No observed significant difference in the lipid-positive proportion per area. AKO animals do not show significant difference in hepatic cholesterol **(c)** and triglycerides **(d)** levels. **(e)** Representative images (40X magnification) from liver histopathology (Masson’s Trichrome - MT staining) from animals exposed to HFD for 28 weeks. The AKO liver samples show a high deposition of collagen compared to WT controls, indicated by the black arrows. **(f)** Quantification of MT-positive cells corroborating the phenotype using Qupath software using the plugin pixel classification by machine learning. **(g)** AKO animals demonstrate a tendency to consume more HFD than WT animals. Data from at least two experiments, each containing at least N=3 mice per genotype. Statistical analyses: Outliers ROUT, Shapiro-Wilk normality test - Unpaired Student t-test. The error bars represent the standard error of the mean (SEM). ns, non-significant (p>0.05); **p<0.01.

**Supplemental Figure 5.**
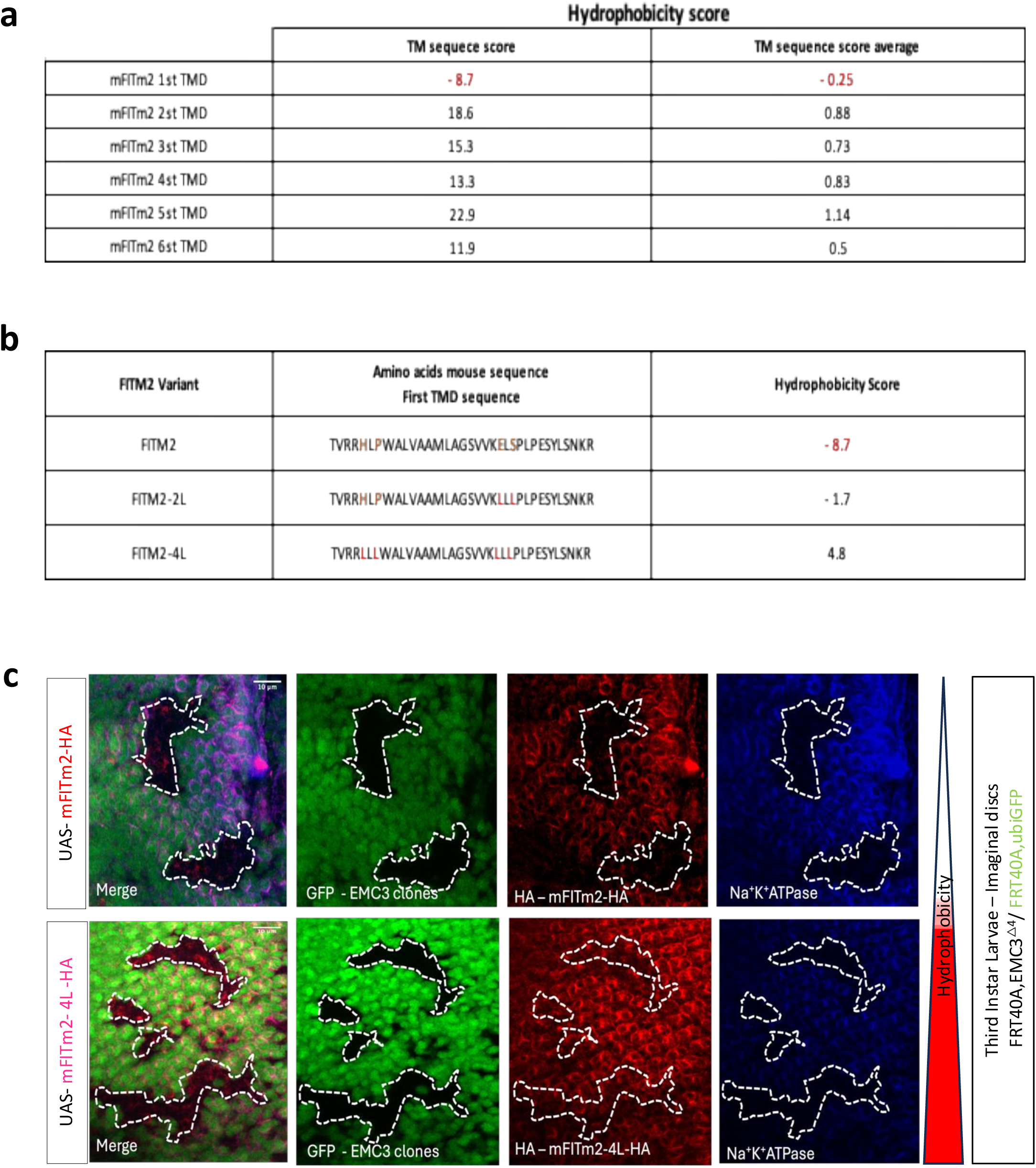
Analysis of FITM2 TMDs hydrophobicity. **(a)** Hydrophobicity analyses of the TMDs of FITM2. The amino acid sequences of each TMD of *Fitm2* were analyzed following the methods described by Zhao & London ^56^. The “transmembrane tendency score” was employed to define the most suitable TMD target for generating the more hydrophobic variants. The first TMD show was identified as being less hydrophobic and was consequently the candidate to be modified. **(b)** Table indicating the point mutations introduced into the first TMD of FITM2. These substitutions, highlighted in red, aim to enhance hydrophobicity and result in FITM2-2L and FITM2-4L mutants. The amino acid hydrophobicity score was calculated following the methods described by Zhao & London ^56^. Each sequence is accompanied by its corresponding “transmembrane tendency score”. **(c)** Representative confocal images of third instar larval eye imaginal discs with eyeless-flippase-induced clones of cells homozygous for EMC3Δ4, denoted by the absence of GFP expression. Territories expressing both the UAS-mFITM2-HA (upper panel) and EMC-independent UAS-mFITM2-4L-HA (lower panel) exhibited reduced expression of the established EMC client Na+K+ATPase α subunit.

## Methods

### Mouse models

An *Emc3* conditional allele *(Emc3 ^lox/+^*), in which exons 4 and 5 of *Emc3* were flanked by LoxP sites, was kindly provided by Jeffrey A.Whitsett ^19^. These *Emc3 ^lox/+^* mice were intercrossed to generate floxed animals (*Emc3^lox/lox^* otherwise denoted *Emc3^floxed^*). *Emc3^floxed^* female mice were mated to Albumin-Cre male *mice (B*6. Cg-*Speer6-ps1^Tg(Alb-cre)21Mgn^*/J - Jackson laboratory – 003574 – Congenic, Transgenic) to generate a conditional deletion of *Emc3* in the liver (*Alb*^cre^ *Emc3*^△/△^) and, mated to Adiponectin-Cre male mice *(B*6; FVB-Tg (Adipoq-cre) 1Evdr/J - Jackson laboratory – 010803 – Transgenic) to generate a conditional deletion of *Emc3* in the white and brown adipose tissues (*Adipoq*^cre^ *Emc3* ^△/△^). The resulting *Alb*^cre^ *Emc3*^△/△^ males and *Adipoq*^cre^ *Emc3*^△/△^ males were mated to *Emc ^floxed^* females. Hence, the offspring were approximately 50% *Emc3^floxed^* (WT) / 50% *Alb*^cre^ *Emc3*^△/△.^ (LKO) and approximately 50% *Emc3^floxed^* (WT) / 50% *Adipoq*^cre^ *Emc3*^△/△.^ (AKO). The floxed littermates was used as a control. All experiments with mice were approved by the Portuguese National Regulatory Agency (DGAV – Direção Geral de Alimentação e Veterinária), the Animal Ethics Committee (ORBEA), and the animal welfare committees of Instituto Gulbenkian de Ciência (Project number A001/2020), following international guidelines for animal care.

### Mouse accommodation and manipulation

The male animals were co-housed after weaning (3 weeks old), to homogenize differences in microbiota and other environmental conditions, in an enriched environment (corn cob bedding and cotton), in acrylic cages with a ventilated air system and an average of 50% humidity (Alesco; six mice/cage), under a controlled dark/light cycle (12/12 h) at standard sub-thermoneutral conditions of 20-24°C at an SPF facility at Instituto Gulbenkian de Ciência. Water and Standard Diet (SD) food (Labina autoclaved mouse chow, Brazil) were provided *ad libitum*. The endpoint of the experiments was when the animals were 8, 12, and 34 weeks old, depending on the protocol. The animals were euthanized with CO_2_ inhalation. For the evaluation of the impact of fasting conditions, the animals were fasted for 12 hours (overnight - O/N) (water *ad libitum*). The animals were monitored once a week, taking into consideration any signs of discomfort or distress and evaluation of food intake and alterations in their body morphology and weight. In the endpoint (depending on each protocol), animals were sacrificed, and total blood samples were collected via cardiac puncture and organ/tissue weight was evaluated. Before tissue harvesting, the animals were perfused with 1X PBS, through cardiac puncture. The liver, interscapular brown adipose tissue (iBAT), inguinal (iWAT), epididymal (eWAT) retroperitoneal (rWAT/RAT), and mesenteric white adipose tissue (MAT) were collected, weighed and processed (frozen in liquid nitrogen and stored at −80°C (embedded in formalin or used as fresh samples). The % tissue weight values were obtained using the formula ((tissue weight (g)/body weight(g))*100)).

### Diet-induced obesity

To induce obesity, 3 weeks old male mice were fed with High Fat Diet (HFD-60 kJ% Fat [Lard] BLUE HF Diet 10 mm, sterilized and irradiated - E15742-347 – SSNIFF Spezial Diet – GMBH), at least, for 9 weeks (or in longer-term experiments for 28 weeks).

### ALT and AST (serology analysis)

The serum concentrations of Alanine Aminotransferase (ALT) and Aspartate Aminotransferase (AST) were measured to assess signs of liver damage. The total blood from fed animals was harvested from the tail and in, centrifuged at 4600 g, 10 min, R.T. The serum samples were collected and sent to DNAtech – Laboratório de Análises Clínicas Veterinárias (Portugal: https://dnatech.pt/), to be analysed.

### Cardiogreen clearance rate assay

Cardiogreen was administrated via intravenous (i.v.) injection at a dosage of 20 mg/kg. After 20 min, total blood was harvested (via puncture of the inferior vena cava in the liver) and centrifuged at 4600 g, 10 min, RT. The serum samples were diluted and aliquoted into a 96-well polystyrene plate (SARSTEDT - 82.1581) and the absorbance was measured using a microplate reader (Thermo Scientific Multiskan Go Cat. N° N10588) at a wavelength of 800 nm, as previously described (Alvarenga *et al.*, 2018). The concentration was quantified based on a Cardiogreen standard curve.

### Histopathology analysis

The liver and adipose tissues from mice were conserved in Formaldehyde 4% Formalin10% buffered (VWR) solution and processed to paraffin-embedded sections (3 µm section) and stained. Liver tissue slides were subsequently stained with various dyes: H&E (Hematoxylin and Eosin stain) and Masson’s Trichrome stain (fibrosis detection). An immunohistology protocol was used to detect cell death and cell proliferation, respectively, using Caspase-3 and KI-67 antibodies at the concentration of 1:200. Liver tissue slides were evaluated for Oil Red-O staining in an 8-um liver cryo-sectioned from liver samples dehydrated in a 30% sucrose; 0,1% sodium azide solution after 48 hours in buffered 10% formalin solution and snap frozen in liquid nitrogen embedded in OCT. The slides containing the sections were scanned into digital images (NanoZoomer-SQ Digital slide scanner – Hamamatsu). 10X liver images were taken and analyzed using Image J software. The lipid droplet’s area was measured using a color threshold plugin, and the fraction within the liver area was estimated. iBAT and eWAT/iWAT were processed to paraffin-embedded sections (3 μm sections), stained with H&E, and scanned into digital images (NanoZoomer-SQ Digital slide scanner -Hamamatsu). The average eWAT and iWAT adipocyte size in adipose tissue sections [expressed as the average cross-sectional area per cell (μm2)] was determined using Fiji software, based on a method described in Parlee et al. (2014). An average of 11000 lipid droplets per mouse were measured. Quantifying size and number of adipocytes in adipose tissue. All histopathological slides were analyzed in a blinded manner by a pathologist. All histopathological slides were analyzed in a blinded manner by a pathologist.

### Transmission Electron Microscopy (TEM)

Overnight fasted animals were euthanatized and perfused through cardiac puncture primary with a 1% PBS solution followed by perfusion with a fixative buffer (2% (v/v) formaldehyde (Science Services EMS, Cat#15710), 2.5% (v/v) glutaraldehyde (Science Services EMS, Cat #16220) in 0.1M Phosphate buffer (PB)). Liver samples were dissected, cut into small pieces (maximum size of 1mm^3^), and immersed for 1h in the fixative buffer under shaking. After the initial fixation, all processing steps, except washes and embedding, were done using a PELCO Biowave Pro+ microwave system. The samples were washed three times with 0.1M PB and post-fixed in 1% (w/v) Osmium tetroxide (Science Services EMS, Cat #19110), 1% (w/v) Potassium Ferrocyanide (Sigma-Aldrich P3289) in 0.1M PB for seven 2 minutes intercalated cycles at 0 or 100 W in vacuum. Then, the samples were washed twice with a 0.1M PB followed by a wash step with distilled water (twice) and stained with 1% (w/v) Tannic Acid (Science Services EMS, Cat #21700) for seven 1-minute intercalated cycles at 0 or 150 W in vacuum. Next, samples were washed with distilled water, five times, for five minutes each. The *En-bloc* staining was performed with a 0.5% (w/v) aqueous solution of uranyl acetate (AnalaR, Cat #10288) for seven 1-minute intercalated cycles at 0 or 150 W in vacuum. An ethanol serial dehydration (30%, 50%, 75%, and 90% was performed for 40 seconds at 150 W for each step. The final dehydration step was three times 100% ethanol for 40 seconds each at 150 W. An infiltration with EMbed-812 epoxy resin (Science Services EMS, Cat #19000) in an increased concentration step (25%, 50%, 75% in ethanol, followed by 100% resin) was realized using a cycle of 3 minutes at 250 W per step. A final step of inclusion in 100% EMbed-812 epoxy resin with DMP-30 was performed to embed samples. The resin was cured at 60°C for at least 24 hours. A Leica UC7 Ultramicrotome was used to obtain 70 nm sections that were collected on palladium-copper grids coated with 1% (w/v) formvar (Agar Scientific, Cat #1201) in chloroform (VWR, Cat#22706.292). Uranyl acetate, in a 1% concentration (w/v)(Agar Scientific, Cat #10288) and Reynolds’ Lead Citrate were used to post-stain the sections (for 5 min each). The sections were imaged on an FEI Tecnai G2 Spirit BioTWIN (120 kV) transmission electron microscope using an Olympus-SIS Veleta CCD Camera.

### Protein extraction quantification and normalization

Mouse liver tissue was collected and lysed using lysis buffer (1% Triton X-100, 150 mM NaCl, 50 mM Tris-HCl, pH 7.4, 5 mM EDTA) supplemented with protease inhibitors (Roche). Samples were disrupted in a TissueLyserII at the highest frequency (30) for 4 minutes and were subsequently centrifuged for 30 min, 21,130 g at 4°C. The supernatant was collected. Mouse adipose tissue was collected and lysed using RIPA lysis buffer (50 mmol/L Tris-HCl (pH 8.0), 0.25 mol/L NaCl, 5 mmol/L EDTA) supplemented with protease inhibitors, and 10 mM 1,10 phenanthroline. The tissue homogenization was done in a TissueLyserII under the highest frequency (30) for 4 min. The homogenate was centrifuged at 6000 g for 15 min, at 4°C. The supernatant was collected. Triton X-100 was added to a final concentration of 1% (v/v), mixed, and incubated at 4°C for 1h.To remove the upper layer of lipids, the samples were centrifuged at 12000 g for 15 min at 4°C and the supernatants were collected in a new Eppendorf. This step was repeated twice to ensure maximal lipid removal. The protein quantification and normalization were performed following the Bradford protocol (DC protein Assay BIORAD) against a BSA concentration curve which was analyzed to fit linear regression (y=mx+b) to obtain the coefficient of determination (R^2^). Equal amounts of protein (approximately 30 µg per well) from different samples were solubilized in Laemmli sample buffer (LDS buffer - supplemented with DTT – final concentration 1X) and denatured by heating at 65°C for 15 min. More details of the reagents and equipments used are provided in Supplementary Table 1 and 4.

### Plasmids

Human FITM2 cDNA was obtained from the mammalian genome collection^66^, amplified by PCR using the primers (FITM2-F: 5’-gatccACCGGTgagcatctggagcgctg-3’ and FITM2-R: 5’-tctgGCGGCCGCttatttcttgtaactatcttgcttc-3’). Both PCR product and a pcDNA5/FRT/TO vector (Invitrogen) already encoding an HA tag at the 5’ end, were digested by AgeI and NotI and resulting products isolated and ligated to yield a plasmid that produces an N-term HA tag (YPYDVPDYA) in frame with FITM2 (HA-FITM2).

#### Compounds and chemicals

The following compounds were used in experiments: Doxycycline (DOX, Sigma-Aldrich), MG132 (Merck Chemicals Ltd), CB-5083 (kind gift of Cleave Biosciences), Bortezomib (BTZ, Merck Chemicals Ltd.)

### Cell culture

U2OS Flp-In^TM^ TRex^TM^ cell lines used in assays have been described in detail previously (^9,10^ and include wild-type (WT), ΔEMC6 (generated by CRISPR-Cas9) and ΔEMC6+rescue (stably expressing FLAG-HA-EMC6 under a Tet-inducible promoter). All cell lines were maintained at 5% CO_2_ and 37°C in Gibco® GlutaMAX® Dulbecco’s Modified Eagle Medium (DMEM; Life Technologies) containing 10% v/v foetal calf serum (FCS) and 2 mM L-glutamine (Life Technologies) (DMEM/10% FCS). Cell lines stably expressing HA-FITM2-pcDNA5 were generated by Flp recombinase-mediated integration and were continuously cultured in DMEM/10% FCS + 100 µg/ml hygromycin B (Hygromycin B Gold, InvivoGen).

### Western blotting

Lysate samples (tissue or cell) were loaded onto an SDS-PAGE (12 or 14%) gel in an electrophoresis chamber containing 1X MOPS Running Buffer (50 mM of MOPS, 50 mM of Tris base, 3.47 mM of SDS, and 1 mM of EDTA). After electrophoresis, the proteins were transferred onto the PVDF membrane in a 1X cold Transfer Buffer (25 mM of Tris, 192 mM of Glycine, and 10 % of Methanol) solution, for 1 h. The membranes were blocked in 1X TBS-T (10 mM Tris pH 7.4, 0.3 M NaCl, 1% Tween) supplemented with 5% non-fat dried milk for 1 h. The 1X TBS-T, 5% milk solution was used to dilute the primary and secondary antibodies. Membranes were incubated with primary antibody overnight at 4°C. Subsequently, the membranes were washed in 1X TBS-T for 10 min, 4 times, and incubated with the appropriate secondary antibody at RT, for 2 hrs. Membranes were then washed, following the sequence: 1X TBS-T, 10 min, 3 times; 1X PBS, 5 min, twice. ECL was used to detect the protein bands, using the Amersham^TM^ imager 680. The images were captured and analyzed under Fiji (ImageJ) software. Western blotting quantification was based on ROI selection and intensity measurement. Details of the antibodies used are provided in Supplementary Table 2.

### Neutral lipid extraction, cholesterol and triglycerides levels evaluation

Fresh liver samples were weighed (50-100 mg/each). A mixture of Chloroform: Methanol (2:1) was added to the tissue in an Eppendorf containing metal spheres for mechanical disruption. The tissue samples were exposed to the highest frequency (30) for 2 min in a TissueLyserII to promote tissue disruption. The lysates were transferred to a glass screw-capped tube, supplemented with 3 mL of Chloroform: Methanol (2:1) and O/N incubated under shaking at RT. On the next day, samples were centrifuged at 800 g, 15 min, RT. The supernatants (3 mL) were transferred to a clean tube. 0.6 mL of 0.9% NaCl was added to each sample (1/5 of the total initial volume), which was then briefly vortexed. 0.5 mL of the lower chloroform phase was transferred to a new glass tube and left at the fume hood to dry (2-3 days) to evaporate. Dried pellets were diluted in 0.5 mL of a mixture of Butanol/(Triton-X-100:Methanol (2:1)) (30:20) and analysed for triglyceride and cholesterol levels using the Kit Spinreact Enzymatic Colorimetric. By performing 1:2 serial dilutions, a calibration curve was prepared (Cal Std stock 200 mg/dL) starting at 100 mg/dL to 0.78 mg/dL in an appropriate solvent solution, depending on the processed samples (0,9% NaCl – liver or serum). 15 µL of standards were loaded in duplicate into a 96-well plate alongside the standards and blanks. 135 µL of working reagent (WR) were subsequently added to each sample. For cholesterol quantification, 5 µL of serum and feces and 15 µL of neutral lipids from the liver were loaded. For the quantification of triglycerides, 5 µL of serum and neutral lipids from the liver were loaded, always in duplicate. The WR was added to the final volume of 150 µL. Samples were incubated at 37°C for 5 min and the absorbance was evaluated in a plate spectrophotometer at 505 nm. Results normalized by standard sample and expressed in mg/dL. The liver values were normalized by liver mass used (mg/g).

### Free glycerol content from serum samples

Blood samples were collected from the tail vein from animals fed with SD and HFD, after O/N fasting. A centrifugation step (7000 rpm/10 min/4°C) was applied to collect the serum samples. Free glycerol reagent (Sigma) and Glycerol Standard Solution (Sigma) were used following the manufacturer’s instructions. Samples were analyses at the absorbance 540 nm and the glycerol concentration were obtained through calculation (A sample – A blank) / (A Standard – A blank) X Concentration of standard (0,26 mg/mL).

### Quantitative Transcriptional Analysis

Mouse liver and iBAT samples were snap frozen a lysed in NR buffer of the NZY Total RNA isolation kit (Nzytech). The total RNA isolated was purified following the protocol present in the same kit and it was used as a template for first-strand cDNA synthesis using an Xpert cDNA Synthesis Supermix (GRiSP). Quantitative PCR was realized with Xpert Fast SYBR (GRiSP) in the ABI QuantStudio 7 system. The protocols were run according to the manufacturer’s instructions. To compare the relative expression of a target gene, the delta Ct value was obtained by applying the following formula: ΔCt = (Ct target gene – Ct of the average of reference genes). More than a single reference gene is recommended by the Guidelines (Bustin *et al.*, 2009) as a good practice for gene expression normalization. To calculate the fold expression the 2^DDCt^ method was used (Schmittgen and Livak, 2008). The primer sequences used are described in Supplementary Table 3.

### Mass Spectrometry and Proteomic Analysis

#### Tissue lysate

Mouse tissues (liver and adipose tissue) were lysed in buffer containing 4% SDS, 100 mM Tris-Cl (pH 7.2), 150 mM NaCl, 5% glycerol, and 1x protease inhibitor cocktail. Zirconium beads were added to each tube and samples subjected to 3 rounds of 30 s bead beating. Lysate was then transferred to a new tube and cellular debris pelleted by centrifugation (10000 g, 10 min). Supernatants including the lipid layer were then transferred to a new tube. Chloroform methanol protein extraction was carried out to isolate proteins. To 200 µL lysate, 600 µL methanol was added followed by 150 µL chloroform, then 450 µL MilliQ water with vortexing after each addition. Samples were then centrifuged (14000 x g, 2 min) and the top aqueous layer was removed. An additional 450 µL aliquot of methanol was added to the remaining sample and vortexed. Samples were centrifuged (14000 x g, 3 min), after which the supernatant was removed and the protein pellet allowed to air dry. Proteins were resuspended in 5% SDS, 100 mM Tris-Cl by first shaking at room temperature (RT) for 30 min and then passing through a syringe. Resolubilized proteins were then quantified using the Pierce™ BCA Protein Assay Kit using the manufacturer’s instructions. Protein levels were normalized to 1 µg/µL in 5% SDS,100mM Tris-Cl buffer. Samples were then reduced by addition of DTT to a final concentration of 20 mM and incubated at room temperature for 30 min. Free thiols were then alkylated by adding Iodoacetamide to a final concentration of 40 mM and incubated in the dark at RT for 30 min. Proteins were then acidified by addition of phosphoric acid to a final concentration of 1.2%. Proteins were then cleaned using 100 µg S-trap column (Protifi). Samples were diluted in S-trap binding buffer (100 mM TEAB in 90% methanol) at a 7:1 binding buffer-to-sample ratio. Proteins were then loaded onto the S-trap column by adding 200 µL of sample to the column and centrifuging (4000 x g, 30 s). This was repeated until the whole sample was loaded. Proteins were then washed by adding 150 µL binding buffer and centrifuging thrice (4000 x g, 30 s). The washed column was then moved to a new tube and trypsin was added to the column at a ratio of 1:25 (trypsin: sample protein) in 30 µL 50 mM TEAB and encouraged into the column with pressure on the top of the tube, then capped and incubated at 47 ⁰C for 2 hours. The resultant peptides were eluted from the column by first adding 40 µL of 50 mM TEAB and centrifuging for 30 sec at 4000 g then adding 40 µL of 0.2% TFA, centrifuging (4000 x g, 30 s), and finally adding 40 µL 50% acetonitrile 0.2% TFA followed by centrifugation (4000 x g, 30 s). The resultant peptide mix was dried down using a rotor vac and resuspended in buffer A (0.2% TFA, 2% acetonitrile, HPLC grade H_2_0).

#### MS downstream data processing

Quantitative outputs from DIA-NN spectra were analysed using the Perseus software platform (v1.6.15.0). Samples were annotated and filtered to have at least 2 valid values. Raw values underwent log2 transformation and were normalized by median subtraction. Volcano plots were then generated to identify proteins that changed significantly between WT and KO tissues. The resulting fold-change and p-values for each protein were exported to Excel for additional processing and filtering. The resulting dataset was filtered further to extract proteins predicted to contain 1 (or more) TMDs and then organized into membrane protein topological classes. The sequences corresponding to first predicted TMD of membrane protein were extracted from SwissProt entries and analysed using the Kyte-Doolittle (K-D) hydrophobicity scale to gauge potential as EMC clients. Briefly, tail anchored (TA) proteins with scores <35 were ranked highest with Rank 1 and those with >35 received Rank 2. For multi-pass proteins, the first TMD was assessed for presence <100 amino acids from the N-terminus and a K-D score <35. Those satisfying both criteria received Rank 1, those satisfying one criterion received Rank 2, and those satisfying neither received Rank 3. Single-pass signal anchor proteins with a TMD within 100 amino acids from the N-terminus and a K-D score <35 received Rank 3 and otherwise, received Rank 4. Proteins containing a signal peptide received Rank 4. Type 3 single-pass TM proteins were also investigated further. A literature and database search of proteins in Rank 1 was then undertaken to identify any phenotypic similarities of KOs in lipid dysregulation (in liver or WAT/BAT) that resembled what was observed for EMC3 KO mice.

### Acute cold challenge

Animals were routinely housed at room temperature (approximately of 22°C) in IGC’s SPF facility. To evaluate the animal’s ability to realize thermoregulation, adipose tissue (*Adipoq*^cre^ *Emc3* ^△/△^) (AKO) and *Emc3 ^floxed^* (WT) male mice were acclimatized in a heat-controlled chamber (Climatic Chamber FITOCLIMA 600 PLHv – Aralab – Serial number 2698) at thermoneutrality (30°C) for one week. Animals were then exposed to acute cold (4°-6°C) and rectal temperature was measured (Thermometer with a rectal probe - AZ8851 K.J.T Type (BioSeB) – Serial number 9191582) respecting the following timing points after exposition: 0.1hr, 2hrs, 3hrs, and 4 hrs.

### Drosophila stocks

The flies were raised at 25°C under a 12-hour light/12-hour dark cycle, in a standard cornmeal. The *Emc3^△4^* mutant allele (*dPob^Δ4^* gene), (Dmel\EMC3 – CG6750 – FlyBase ID FBgn0032292) and Lsp2-Gal4 (III) were kind offers by Akilo Satoh and Alisson Gontijo, respectively.

### Plasmids to generate transgenic flies for expression of HA-tagged mFITm2 and hydrophobic variants

The plasmids construct for overexpression in *Drosophila* of the mouse *fitm2* coding sequence, named here as *mfitm2*, and the mutant variants with increased hydrophobicity in the first TMD (*mFITm2*-2L and *mFITm2-*4L) were generated by amplifying the first TMD and flanking residues of mFITm2. To generate a more hydrophobic first TMD we substituted the following amino acids from the *mfitm2* original mouse coding sequence gene in their sequence corresponding to the first TMD (VRATVRRHLPWALVAAMLAGSVVKELSP) with Leucine (L). For mFITm2 −2L, L was inserted instead of the Glutamic Acid (E) 40 and Serine (S) 42 (VRATVRRHLPWALVAAMLAGSVVKLLLP). For mFITm2 −4L, L was inserted instead of Histidine (H) 23, Proline (P) 25, E 40, and S 42 (VRATVRRLLLWALVAAMLAGSVVKLLLP). After the amplification of the inserts, we proceeded with Gibson Assembly® Cloning (New England Biolabs) using the linearized plasmid pUASTattB (previously treated with EcoRI) and the C-terminal fragment of mFITm2 to make a full-length protein.

Production of HA-tagged mFITm2, mFITm2–2L, and mFITm2–4L transgenic flies were conducted by PCR amplification of the full open reading frame of mFIT2 from the plasmid pMDC32-mFIT2, kindly provided by Prof.Dr. David Silver ^6^. To generate the C-terminally HA-tagged mFITm2-(HA), we used the following primer sequences to amplify by PCR: 5’-GGGAATTCGTTAACAGATCTATGGAGCACCTGGAGCGCT-3’and5’-CGAGCCGCGGCCGCAGATCTTCAAGCGTAATCTGGAACATCGTATGGGTATTTCTTGTAAGTATCTCGC -3’. To create the 2L fly line, a two steps PCR protocol was used: (PCR1) 5’-GGGAATTCGTTAACAGATCTATGGAGCACCTGGAGCGCT-3’and 5’GCTCTCGGGCAGCGGCAGCAGCAGCTTGACGACCGAGCC-3’ ; (PCR2) 5’GGCTCGGTCGTCAAGCTGCTGCTGCCGCTGCCCGAGAGC 3’ and 5’CGAGCCGCGGCCGCAGATCTTCAAGCGTAATCTGGAACATCGTATGGGTA TTTCTTGTAAGTATCTCGC 3’. To generate the 4L version, two PCR amplification reaction were performed using the following primers: (PCR1) 5’-GGGAATTCGTTAACAGATCTATGGAGCACCTGGAGCGCT -3’ and 5’-CATCGCCGCCACCAGCGCCCACAGCAGCAGGCGGCGCACGGTCGCGCGCAC -3’; and (PCR2) 5’-TGCGCGCGACCGTGCGCCGCCTGCTG CTGTGGGCGCTGGTGGCGGCGATG-3’ and 5’-5’CGAGCCGCGGCCGCAGATCTTCAAGCGTAATCTGGAACATCGTATGGGTATTTCTTGTAAGTATCT CGC -3’, using 2L as template DNA The inserts were then amplified by PCR with the primers previously described and assembled using Gibson assembly to produce C-terminal HA-tagged pUAST.attB constructs. DNA purification was procedure using Miniprep or MidiPrep (Nzytech). All the purified DNA samples were sent to STAB VIDA to be sequenced. The constructs were injected into *Drosophila* embryos through microinjection into the attB P2 site in the 3rd chromosome through PhiC31 integrase-mediated transgenesis at the Champalimaud Foundation transgenics facility.

### Assays in Drosophila imaginal discs

To assess whether mFITm2 expression depends on the EMC, we performed crosses between fly stocks carrying C-terminally HA-tagged mFITm2 constructs on the third chromosome and flies harbouring an *Emc3^△4^* mutant allele, the *dPob^Δ4^* FRT40A (Dmel\EMC3 – CG6750 – Fly Base ID FBgn0032292)^14^on the second chromosome. The resultant flies (genotype: w; *dPob^Δ4^* FRT40A; UAS.mFITm2-HA, or their more hydrophobic 2L and 4 L variants, were subsequently crossed with eyFLP, GMRGAL4; ubiGFPFRT40A flies to generate EMC3^△4^ mutant mosaic eye imaginal discs expressing mFITm2-HA proteins. The imaginal discs immunostaining and imaging were realized following protocol previously described ^7^. The Zeiss LSM 880 confocal microscope was used to obtain images, under objective 20X, that were analyzed in ImageJ – FIJI (version 2.1.0/1.53C). For antibody information, consult Supplementary Table 2.

### Induction of EMC mutant mosaic tissues in the fat body induced by heat shock

The fat body mosaic tissue was generated by crossing males bearing the *Emc3^△4^* mutant chromosome with female virgins caring a heat shock inducible Flipase (hs-Flp) in the X chromosome and Ubi-GFP FRT in the 2^nd^ chromosome (Males: w/Y;dPobΔ4FRT40A / CyO; Sb/TM6B X hsflp; ubiGFPFRT40A / CyO. The parents were left in vials for 72 hrs before being transferred to a new vial. After 6 hours, the parents were removed and the eggs were submitted to a heat shock (37°C, 1hr), to induce the expression of the Flipase that mediates recombination between the chromosomes containing the FRT sites generating, after segregation of the chromosomes during mitosis, cells that are homozygous for the EMC3^△4^ mutation, labeled by the absence of GFP expression and GFP positive control (non-mutant) cells. To evaluate the mFITm2-4L rescue phenotype, the fat body mosaic tissue was generated crossing males with the genotype hs-Flp; UbiGFP, FRT40A; Lsp2-Gal4 with virgin females with the genotype w; dPobΔ4, FRT40A; UAS-mFITm2-L4-HA. Third instar stage larvae from this cross were selected (hs-Flp; UbiGFP.FRT40A/ dPobΔ4FRT40A; Lsp2-Gal4/UASmFITm2-4L-HA) after heat shock protocol to generate the *Emc3^△4^* mutant clones.

### Fat body immunostaining and imaging

One week after egg laying, third instar stage larvae were collected, and the fat bodies were dissected in 1X PBS, fixed in 4% PFA (paraformaldehyde) at RT for 1 h. A washing step in 1X PBS was performed three times, 10 min each, at RT, under gentle shaking. Lipid droplets were stained using a Nile Red Staining Solution, 10 min of incubation, in the dark, at RT (concentration of 0.25 mg/mL -1 uL diluted in 4 mL of 1X PBS). A rinse step was performed using 1X PBS and immediately, fat body samples were mounted in Vectashield (with DAPI) (Vector Laboratories). Fixation, washing, and staining were performed under shaking. The Zeiss LSM 880 confocal microscope was used to obtain images, under the 20X objective, that were analyzed in ImageJ – FIJI (version 2.1.0/1.53C). DNA was stained with DAPI (in blue), GFP expression was visualized in the green channel and the lipid droplets were observed in the red channel. The significance was evaluated using GraphPad Prism 9 software. For dye information, consult Supplementary Table 2.

## Notes

### Competing Interest Statement

The authors have declared no competing interest.

## References

1. Hegde, R. S. & Keenan, R. J. A unifying model for membrane protein biogenesis. Nat Struct Mol Biol 31, 1009–1017 (2024).

2. Anghel, S. A., McGilvray, P. T., Hegde, R. S. & Keenan, R. J. Identification of Oxa1 Homologs Operating in the Eukaryotic Endoplasmic Reticulum. Cell Rep 21, 3708–3716 (2017).

3. Lewis, A. J. O. & Hegde, R. S. A unified evolutionary origin for the ubiquitous protein transporters SecY and YidC. BMC Biol 19, 266 (2021).

4. Borgese, N. Getting membrane proteins on and off the shuttle bus between the endoplasmic reticulum and the Golgi complex. J Cell Sci 129, 1537–1545 (2016).

5. McGilvray, P. T., Anghel, S. A., Sundaram, A., Zhong, F., Trnka, M. J., Fuller, J. R., Hu, H., Burlingame, A. L. & Keenan, R. J. An ER translocon for multi-pass membrane protein biogenesis. Elife 9, e56889 (2020).

6. Chen, K., Wang, Y., Yang, J., Klöting, N., Liu, C., Dai, J., Jin, S., Chen, L., Liu, S., Liu, Y., Yu, Y., Liu, X., Miao, Q., Liew, C. W., Wang, Y., Dietrich, A., Blüher, M. & Wang, X. EMC10 modulates hepatic ER stress and steatosis in an isoform-specific manner. J Hepatol 81, 479–491 (2024).

7. Page, K. R., Nguyen, V. N., Pleiner, T., Tomaleri, G. P., Wang, M. L., Guna, A., Hazu, M., Wang, T.-Y., Chou, T.-F. & Voorhees, R. M. Role of a holo-insertase complex in the biogenesis of biophysically diverse ER membrane proteins. Mol Cell 84, 3302–3319.e11 (2024).

8. Wu, H., Smalinskaitė, L. & Hegde, R. S. EMC rectifies the topology of multipass membrane proteins. Nat Struct Mol Biol 31, 32–41 (2024).

9. Guna, A., Volkmar, N., Christianson, J. C. & Hegde, R. S. The ER membrane protein complex is a transmembrane domain insertase. Science 359, 470–473 (2018).

10. Volkmar, N., Thezenas, M. L., Louie, S. M., Juszkiewicz, S., Nomura, D. K., Hegde, R. S., Kessler, B. M. & Christianson, J. C. The ER membrane protein complex promotes biogenesis of sterol-related enzymes maintaining cholesterol homeostasis. J Cell Sci 132, jcs223453 (2019).

11. Gaspar, C. J., Vieira, L. C., Santos, C. C., Christianson, J. C., Jakubec, D., Strisovsky, K., Adrain, C. & Domingos, P. M. EMC is required for biogenesis of Xport-A, an essential chaperone of Rhodopsin-1 and the TRP channel. EMBO Rep 23, e53210 (2022).

12. O’Keefe, S., Zong, G., Duah, K. B., Andrews, L. E., Shi, W. Q. & High, S. An alternative pathway for membrane protein biogenesis at the endoplasmic reticulum. Commun Biol 4, 828 (2021).

13. Hiramatsu, N., Tago, T., Satoh, T. & Satoh, A. K. ER membrane protein complex is required for the insertions of late-synthesized transmembrane helices of Rh1 in Drosophila photoreceptors. Mol Biol Cell 30, 2890–2900 (2019).

14. Satoh, T., Ohba, A., Liu, Z., Inagaki, T. & Satoh, A. K. dPob/EMC is essential for biosynthesis of rhodopsin and other multi-pass membrane proteins in Drosophila photoreceptors. Elife 4, e06306 (2015).

15. Xiong, L., Zhang, L., Yang, Y., Li, N., Lai, W., Wang, F., Zhu, X. & Wang, T. ER complex proteins are required for rhodopsin biosynthesis and photoreceptor survival in Drosophila and mice. Cell Death Differ 27, 646–661 (2020).

16. Sun, K., Liu, L., Jiang, X., Wang, H., Wang, L., Yang, Y., Liu, W., Zhang, L., Zhao, X. & Zhu, X. The endoplasmic reticulum membrane protein complex subunit Emc6 is essential for rhodopsin localization and photoreceptor cell survival. Genes Dis 11, 1035–1049 (2024).

17. Zhu, X., Qi, X., Yang, Y., Tian, W., Liu, W., Jiang, Z., Li, S. & Zhu, X. Loss of the ER membrane protein complex subunit Emc3 leads to retinal bipolar cell degeneration in aged mice. PLoS One 15, e0238435 (2020).

18. Gaspar, C. J., Gomes, T., Martins, J. C., Melo, M. N. & Adrain…, C. Xport-A functions as a chaperone by stabilizing the first five transmembrane domains of rhodopsin-1. Iscience (2023).

19. Tang, X., Snowball, J. M., Xu, Y., Na, C. L., Weaver, T. E., Clair, G., Kyle, J. E., Zink, E. M., Ansong, C., Wei, W., Huang, M., Lin, X. & Whitsett, J. A. EMC3 coordinates surfactant protein and lipid homeostasis required for respiration. J Clin Invest 127, 4314–4325 (2017).

20. Huang, M., Yang, L., Jiang, N., Dai, Q., Li, R., Zhou, Z., Zhao, B. & Lin, X. Emc3 maintains intestinal homeostasis by preserving secretory lineages. Mucosal Immunol 14, 873–886 (2021).

21. Liu, L., Mao, S., Chen, K., Dai, J., Jin, S., Chen, L., Wang, Y., Guo, L., Yang, Y., Zhan, C., Xiong, Z., Diao, H., Zhou, Y., Ding, Q. & Wang, X. Membrane-Bound EMC10 Is Required for Sperm Motility via Maintaining the Homeostasis of Cytoplasm Sodium in Sperm. Int J Mol Sci 23, 10069 (2022).

22. Diamantopoulou, A., Sun, Z., Mukai, J., Xu, B., Fenelon, K., Karayiorgou, M. & Gogos, J. A. Loss-of-function mutation in Mirta22/Emc10 rescues specific schizophrenia-related phenotypes in a mouse model of the 22q11.2 deletion. Proc Natl Acad Sci U S A 114, E6127–E6136 (2017).

23. Reboll, M. R., Korf-Klingebiel, M., Klede, S., Polten, F., Brinkmann, E., Reimann, I., Schönfeld, H. J., Bobadilla, M., Faix, J., Kensah, G., Gruh, I., Klintschar, M., Gaestel, M., Niessen, H. W., Pich, A., Bauersachs, J., Gogos, J. A., Wang, Y. & Wollert, K. C. EMC10 (Endoplasmic Reticulum Membrane Protein Complex Subunit 10) Is a Bone Marrow-Derived Angiogenic Growth Factor Promoting Tissue Repair After Myocardial Infarction. Circulation 136, 1809–1823 (2017).

24. Lin, D. L., Inoue, T., Chen, Y. J., Chang, A., Tsai, B. & Tai, A. W. The ER Membrane Protein Complex Promotes Biogenesis of Dengue and Zika Virus Non-structural Multi-pass Transmembrane Proteins to Support Infection. Cell Rep 27, 1666–1674.e4 (2019).

25. Ngo, A. M., Shurtleff, M. J., Popova, K. D., Kulsuptrakul, J., Weissman, J. S. & Puschnik, A. S. The ER membrane protein complex is required to ensure correct topology and stable expression of flavivirus polyproteins. Elife 8, e48469 (2019).

26. Lahiri, S., Chao, J. T., Tavassoli, S., Wong, A. K., Choudhary, V., Young, B. P., Loewen, C. J. & Prinz, W. A. A conserved endoplasmic reticulum membrane protein complex (EMC) facilitates phospholipid transfer from the ER to mitochondria. PLoS Biol 12, e1001969 (2014).

27. Iyer, A., Niemann, M., Serricchio, M., Dewar, C. E., Oeljeklaus, S., Farine, L., Warscheid, B., Schneider, A. & Bütikofer, P. The endoplasmic reticulum membrane protein complex localizes to the mitochondrial - endoplasmic reticulum interface and its subunits modulate phospholipid biosynthesis in Trypanosoma brucei. PLoS Pathog 18, e1009717 (2022).

28. Shurtleff, M. J., Itzhak, D. N., Hussmann, J. A., Schirle Oakdale, N. T., Costa, E. A., Jonikas, M., Weibezahn, J., Popova, K. D., Jan, C. H., Sinitcyn, P., Vembar, S. S., Hernandez, H., Cox, J., Burlingame, A. L., Brodsky, J. L., Frost, A., Borner, G. H. & Weissman, J. S. The ER membrane protein complex interacts cotranslationally to enable biogenesis of multipass membrane proteins. Elife 7, e37018 (2018).

29. Ebrahimi-Fakhari, D., Wahlster, L., Bartz, F., Werenbeck-Ueding, J., Praggastis, M., Zhang, J., Joggerst-Thomalla, B., Theiss, S., Grimm, D., Ory, D. S. & Runz, H. Reduction of TMEM97 increases NPC1 protein levels and restores cholesterol trafficking in Niemann-pick type C1 disease cells. Hum Mol Genet 25, 3588–3599 (2016).

30. Alves-Bezerra, M. & Cohen, D. E. Triglyceride Metabolism in the Liver. Compr Physiol 8, 1–8 (2017).

31. Sakers, A., De Siqueira, M. K., Seale, P. & Villanueva, C. J. Adipose-tissue plasticity in health and disease. Cell 185, 419–446 (2022).

32. Milesi-Hallé, A., Abdel-Rahman, S. M., Brown, A., McCullough, S. S., Letzig, L., Hinson, J. A. & James, L. P. Indocyanine green clearance varies as a function of N-acetylcysteine treatment in a murine model of acetaminophen toxicity. Chem Biol Interact 189, 222–229 (2011).

33. Comerford, S. A., Hinnant, E. A., Chen, Y. & Hammer, R. E. Hepatic ribosomal protein S6 (Rps6) insufficiency results in failed bile duct development and loss of hepatocyte viability; a ribosomopathy-like phenotype that is partially p53-dependent. PLoS Genet 19, e1010595 (2023).

34. Holloway, M. G., Cui, Y., Laz, E. V., Hosui, A., Hennighausen, L. & Waxman, D. J. Loss of sexually dimorphic liver gene expression upon hepatocyte-specific deletion of Stat5a-Stat5b locus. Endocrinology 148, 1977–1986 (2007).

35. Heffner, C. S., Herbert Pratt, C., Babiuk, R. P., Sharma, Y., Rockwood, S. F., Donahue, L. R., Eppig, J. T. & Murray, S. A. Supporting conditional mouse mutagenesis with a comprehensive cre characterization resource. Nat Commun 3, 1218 (2012).

36. Miller-Vedam, L. E., Bräuning, B., Popova, K. D., Schirle Oakdale, N. T., Bonnar, J. L., Prabu, J. R., Boydston, E. A., Sevillano, N., Shurtleff, M. J., Stroud, R. M., Craik, C. S., Schulman, B. A., Frost, A. & Weissman, J. S. Structural and mechanistic basis of the EMC-dependent biogenesis of distinct transmembrane clients. Elife 9, e62611 (2020).

37. Tozawa, R., Ishibashi, S., Osuga, J., Yagyu, H., Oka, T., Chen, Z., Ohashi, K., Perrey, S., Shionoiri, F., Yahagi, N., Harada, K., Gotoda, T., Yazaki, Y. & Yamada, N. Embryonic lethality and defective neural tube closure in mice lacking squalene synthase. J Biol Chem 274, 30843–30848 (1999).

38. Badman, M. K., Pissios, P., Kennedy, A. R., Koukos, G., Flier, J. S. & Maratos-Flier, E. Hepatic fibroblast growth factor 21 is regulated by PPARalpha and is a key mediator of hepatic lipid metabolism in ketotic states. Cell Metab 5, 426–437 (2007).

39. Kersten, S., Seydoux, J., Peters, J. M., Gonzalez, F. J., Desvergne, B. & Wahli, W. Peroxisome proliferator-activated receptor alpha mediates the adaptive response to fasting. J Clin Invest 103, 1489–1498 (1999).

40. Li, Y., Chao, X., Yang, L., Lu, Q., Li, T., Ding, W.-X. & Ni, H.-M. Impaired Fasting-Induced Adaptive Lipid Droplet Biogenesis in Liver-Specific Atg5-Deficient Mouse Liver Is Mediated by Persistent Nuclear Factor-Like 2 Activation. Am J Pathol 188, 1833–1846 (2018).

41. Ge, S. X., Jung, D. & Yao, R. ShinyGO: a graphical gene-set enrichment tool for animals and plants. Bioinformatics 36, 2628–2629 (2020).

42. Kadereit, B., Kumar, P., Wang, W.-J., Miranda, D., Snapp, E. L., Severina, N., Torregroza, I., Evans, T. & Silver, D. L. Evolutionarily conserved gene family important for fat storage. Proceedings of the National Academy of Sciences 105, 94–99 (2008).

43. Gross, D. A., Snapp, E. L. & Silver, D. L. Structural Insights into Triglyceride Storage Mediated by Fat Storage-Inducing Transmembrane (FIT) Protein 2. PLoS ONE 5, e10796 (2010).

44. Becuwe, M., Bond, L. M., Pinto, A. F. M., Boland, S., Mejhert, N., Elliott, S. D., Cicconet, M., Graham, M. M., Liu, X. N., Ilkayeva, O., Saghatelian, A., Walther, T. C. & Farese, R. V. FIT2 is an acyl-coenzyme A diphosphatase crucial for endoplasmic reticulum homeostasis. J Cell Biol 219, e202006111 (2020).

45. Choudhary, V., Ojha, N., Golden, A. & Prinz, W. A. A conserved family of proteins facilitates nascent lipid droplet budding from the ER. J Cell Biol 211, 261–271 (2015).

46. Seco, C. Z., Castells-Nobau, A., Joo, S.-h., Schraders, M., Foo, J. N., van der Voet, M., Velan, S. S., Nijhof, B., Oostrik, J., de Vrieze, E., Katana, R., Mansoor, A., Huynen, M., Szklarczyk, R., Oti, M., Tranebjærg, L., van Wijk, E., Scheffer-de Gooyert, J. M., Siddique, S., Baets, J., de Jonghe, P., Kazmi, S. A. R., Sadananthan, S. A., van de Warrenburg, B. P., Khor, C. C., Göpfert, M. C., Qamar, R., Schenck, A., Kremer, H. & Siddiqi, S. A homozygousFITM2mutation causes a deafness-dystonia syndrome with motor regression and signs of ichthyosis and sensory neuropathy. Disease Models & Mechanisms (2016).

47. Miranda, D. A., Koves, T. R., Gross, D. A., Chadt, A., Al-Hasani, H., Cline, G. W., Schwartz, G. J., Muoio, D. M. & Silver, D. L. Re-patterning of skeletal muscle energy metabolism by fat storage-inducing transmembrane protein 2. J Biol Chem 286, 42188–42199 (2011).

48. Gross, D. A., Zhan, C. & Silver, D. L. Direct binding of triglyceride to fat storage-inducing transmembrane proteins 1 and 2 is important for lipid droplet formation. Proc Natl Acad Sci U S A 108, 19581–19586 (2011).

49. Cannon, B. & Nedergaard, J. Brown adipose tissue: function and physiological significance. Physiol Rev 84, 277–359 (2004).

50. Eguchi, J., Wang, X., Yu, S., Kershaw, E. E., Chiu, P. C., Dushay, J., Estall, J. L., Klein, U., Maratos-Flier, E. & Rosen, E. D. Transcriptional control of adipose lipid handling by IRF4. Cell Metab 13, 249–259 (2011).

51. Miranda, D. A., Kim, J.-H., Nguyen, L. N., Cheng, W., Tan, B. C., Goh, V. J., Tan, J. S. Y., Yaligar, J., KN, B. P., Velan, S. S., Wang, H. & Silver, D. L. Fat Storage-inducing Transmembrane Protein 2 Is Required for Normal Fat Storage in Adipose Tissue. Journal of Biological Chemistry 289, 9560–9572 (2014).

52. Amin, A., Badenes, M., Tüshaus, J., de Carvalho, É., Burbridge, E., Faísca, P., Trávníčková, K., Barros, A., Carobbio, S., Domingos, P. M., Vidal-Puig, A., Moita, L. F., Maguire, S., Stříšovský, K., Ortega, F. J., Fernández-Real, J. M., Lichtenthaler, S. F. & Adrain, C. Semaphorin 4B is an ADAM17-cleaved adipokine that inhibits adipocyte differentiation and thermogenesis. Mol Metab 73, 101731 (2023).

53. Agrawal, M., Yeo, C. R., Shabbir, A., Chhay, V., Silver, D. L., Magkos, F., Vidal-Puig, A. & Toh, S. A. Fat storage-inducing transmembrane protein 2 (FIT2) is less abundant in type 2 diabetes, and regulates triglyceride accumulation and insulin sensitivity in adipocytes. FASEB J 33, 430–440 (2019).

54. Nishihama, N., Nagayama, T., Makino, S. & Koishi, R. Mice lacking fat storage-inducing transmembrane protein 2 show improved profiles upon pressure overload-induced heart failure. Heliyon 5, e01292 (2019).

55. Chitraju, C., Fischer, A. W., Farese, R. V. & Walther, T. C. Lipid Droplets in Brown Adipose Tissue Are Dispensable for Cold-Induced Thermogenesis. Cell Rep 33, 108348 (2020).

56. Zhao, G. & London, E. An amino acid “transmembrane tendency” scale that approaches the theoretical limit to accuracy for prediction of transmembrane helices: relationship to biological hydrophobicity. Protein Sci 15, 1987–2001 (2006).

57. Musselman, L. P. & Kühnlein, R. P. Drosophila as a model to study obesity and metabolic disease. J Exp Biol 221, jeb163881 (2018).

58. Wang, H., Nikain, C., Amengual, J., La Forest, M., Yu, Y., Wang, M. C., Watts, R., Lehner, R., Qiu, Y., Cai, M., Kurland, I. J., Goldberg, I. J., Rajan, S., Hussain, M. M., Brodsky, J. L. & Fisher, E. A. FITM2 deficiency results in ER lipid accumulation, ER stress, reduced apolipoprotein B lipidation, and VLDL triglyceride secretion in vitro and in mouse liver. bioRxiv 2023.12.05.570183 (2023).

59. Bond, L. M., Ibrahim, A., Lai, Z. W., Walzem, R. L., Bronson, R. T., Ilkayeva, O. R., Walther, T. C. & Farese, R. V. Fitm2 is required for ER homeostasis and normal function of murine liver. Journal of Biological Chemistry 299, 103022 (2023).

60. Nagashima, S., Yagyu, H., Tozawa, R., Tazoe, F., Takahashi, M., Kitamine, T., Yamamuro, D., Sakai, K., Sekiya, M., Okazaki, H., Osuga, J., Honda, A. & Ishibashi, S. Plasma cholesterol-lowering and transient liver dysfunction in mice lacking squalene synthase in the liver. J Lipid Res 56, 998–1005 (2015).

61. Wang, X., Li, Y., Qiang, G., Wang, K., Dai, J., McCann, M., Munoz, M. D., Gil, V., Yu, Y., Li, S., Yang, Z., Xu, S., Cordoba-Chacon, J., De Jesus, D. F., Sun, B., Chen, K., Wang, Y., Liu, X., Miao, Q., Zhou, L., Hu, R., Ding, Q., Kulkarni, R. N., Gao, D., Blüher, M. & Liew, C. W. Secreted EMC10 is upregulated in human obesity and its neutralizing antibody prevents diet-induced obesity in mice. Nat Commun 13, 7323 (2022).

62. Leznicki, P., Schneider, H. O., Harvey, J. V., Shi, W. Q. & High, S. Co-translational biogenesis of lipid droplet integral membrane proteins. J Cell Sci 135, jcs259220 (2022).

63. Chen, F., Yan, B., Ren, J., Lyu, R., Wu, Y., Guo, Y., Li, D., Zhang, H. & Hu, J. FIT2 organizes lipid droplet biogenesis with ER tubule-forming proteins and septins. J Cell Biol 220, e201907183 (2021).

64. Hayes, M., Choudhary, V., Ojha, N., Shin, J. J., Han, G. S., Carman, G. M., Loewen, C. J., Prinz, W. A. & Levine, T. Fat storage-inducing transmembrane (FIT or FITM) proteins are related to lipid phosphatase/phosphotransferase enzymes. Microb Cell 5, 88–103 (2017).

65. Benjamini, Y., Krieger, A. M. & Yekutieli, D. Adaptive linear step-up procedures that control the false discovery rate. Biometrika (2006).

66. Gerhard, D. S., Wagner, L., Feingold, E. A., Shenmen, C. M., Grouse, L. H., Schuler, G., Klein, S. L., Old, S., Rasooly, R., Good, P., Guyer, M., Peck, A. M., Derge, J. G., Lipman, D., Collins, F. S., Jang, W., Sherry, S., Feolo, M., Misquitta, L., Lee, E., Rotmistrovsky, K., Greenhut, S. F., Schaefer, C. F., Buetow, K., Bonner, T. I., Haussler, D., Kent, J., Kiekhaus, M., Furey, T., Brent, M., Prange, C., Schreiber, K., Shapiro, N., Bhat, N. K., Hopkins, R. F., Hsie, F., Driscoll, T., Soares, M. B., Casavant, T. L., Scheetz, T. E., Brownstein, M. J., Usdin, T. B., Toshiyuki, S., Carninci, P., Piao, Y., Dudekula, D. B., Ko, M. S. H., Kawakami, K., Suzuki, Y., Sugano, S., Gruber, C. E., Smith, M. R., Simmons, B., Moore, T., Waterman, R., Johnson, S. L., Ruan, Y., Wei, C. L., Mathavan, S., Gunaratne, P. H., Wu, J., Garcia, A. M., Hulyk, S. W., Fuh, E., Yuan, Y., Sneed, A., Kowis, C., Hodgson, A., Muzny, D. M., McPherson, J., Gibbs, R. A., Fahey, J., Helton, E., Ketteman, M., Madan, A., Rodrigues, S., Sanchez, A., Whiting, M., Madari, A., Young, A. C., Wetherby, K. D., Granite, S. J., Kwong, P. N., Brinkley, C. P., Pearson, R. L., Bouffard, G. G., Blakesly, R. W., Green, E. D., Dickson, M. C., Rodriguez, A. C., Grimwood, J., Schmutz, J., Myers, R. M., Butterfield, Y. S. N., Griffith, M., Griffith, O. L., Krzywinski, M. I., Liao, N., Morin, R., Palmquist, D., Petrescu, A. S., Skalska, U., Smailus, D. E., Stott, J. M., Schnerch, A., Schein, J. E., Jones, S. J. M., Holt, R. A., Baross, A., Marra, M. A., Clifton, S., Makowski, K. A., Bosak, S., Malek, J. & MGC Project Team. The status, quality, and expansion of the NIH full-length cDNA project: the Mammalian Gene Collection (MGC). Genome Res 14, 2121–2127 (2004).

